# 3DVascNet: an automated software for segmentation and quantification of vascular networks in 3D

**DOI:** 10.1101/2023.10.19.563201

**Authors:** Hemaxi Narotamo, Margarida Silveira, Cláudio A. Franco

## Abstract

**Background:** Analysis of vascular networks is an essential step to unravel the mechanisms regulating the physiological and pathological organization of blood vessels. So far, most of the analyses are performed using 2D projections of 3D networks, a strategy that has several obvious shortcomings. For instance, it does not capture the true geometry of the vasculature, and generates artifacts on vessel connectivity. These limitations are accepted in the field because manual analysis of 3D vascular networks is a laborious and complex process that is often prohibitive for large volumes.

**Methods:** To overcome these issues, we developed 3DVascNet, a deep learning (DL) based software for automated segmentation and quantification of 3D retinal vascular networks. 3DVascNet performs segmentation based on a DL model, and it quantifies vascular morphometric parameters such as the vessel density, branch length, vessel radius, and branching point density.

**Results:** We tested 3DVascNet’s performance using a large dataset of 3D microscopy images of mouse retinal blood vessels. We demonstrated that 3DVascNet efficiently segments vascular networks in 3D, and that vascular morphometric parameters capture phenotypes detected by using manual segmentation and quantification in 2D. In addition, we showed that, despite being trained on retinal images, 3DVascNet has high generalization capability and successfully segments images originating from other datasets and organs. Moreover, the source code of 3DVascNet is publicly available, thus it can be easily extended for the analysis of other 3D vascular networks by other users.

**Conclusions:** Overall, we present 3DVascNet, a freely-available software that includes a user-friendly graphical interface for researchers with no programming experience, which will greatly facilitate the ability to study vascular networks in 3D in health and disease.

## 1 Introduction

Blood vessels organisation and function are essential for embryogenesis and homeostasis, and their dysregulation is associated with several diseases, including cancer, cardiovascular diseases, neurodegenerative disorders and ageing. Blood vessels form an intricate and complex network of vessels that irrigate all tissues. The three-dimensional (3D) hierarchical architecture of vessels is relevant for their correct function, and is stereotypical to each organ [1–3]. Yet, quantitative information on vascular parameters is mostly extracted from maximum-intensity projections (MIP) of stacks of images, that is, quantifications are performed in two-dimensional (2D) images [4–9].

Researchers have adopted this conscious limitation because manual analysis of 3D vascular networks is a laborious and complex process that is often prohibitive for large volumes. However, 2D analysis brings obvious shortcomings. For instance, analyses based on 2D MIPs from 3D images neglect the 3D anatomical structure of the vasculature, thereby missing important 3D biological features and not capturing the full complexity of phenotypes involved in physiology and pathology. Recently, researchers have started to use deep learning (DL) methods with automated and semi-automated image processing pipelines to segment vascular networks in 3D. For instance, Todorov et al. [10] presented VesSAP, an automated tool for 3D segmentation and quantification of the mouse brain vasculature. In VesSAP, segmentation is performed using a 3D convolutional neural network. Nevertheless, this method and most of the DL models that have been developed for vessel segmentation require manually annotated datasets for training [11–13]. However, manual segmentation is a time-consuming process and, particularly for 3D blood vessels, it is often prohibitive, which hinders significantly the development of such methods.

To overcome these limitations, we have developed 3DVascNet, a DL-based software for automated segmentation and quantification of 3D vascular networks, that does not require pairs of images and manually-annotated ground-truth segmentation masks. To develop 3DVascNet, we focused on retinal blood vessels. Dysfunction of this stereotypical vascular network is associated with ocular diseases, such as diabetic retinopathy, age-related macular degeneration, and choroidal neovascularization [11, 14], and it is one of the reference models to investigate angiogenesis [14]. Hence, over the years, significant attention has been paid to retinal blood vessel formation and remodelling [11, 14].

Several alterations in the morphology of the blood vessels provide valuable information for the diagnosis of retinal diseases. Various geometric features are typically used to describe the structure and morphology of the blood vessels: diameter, length, area, perimeter, tortuosity, number of branches, and branching points and angles [13, 15]. On the one hand, for clinicians, the analysis and interpretation of these features are important for the diagnosis of several pathological conditions [11–13]. On the other hand, for researchers, the vessels’ features are relevant to understand the mechanisms underlying physiological and pathological retinal vasculature formation [14], which is fundamental to develop new therapeutic approaches [16].

The proposed 3DVascNet software enables the analysis of blood vessel networks in 3D, which will greatly facilitate the ability to quantify and study vascular networks in health and disease. 3DVascNet is based on a CycleGAN model [17], a generative DL model designed to translate images from a domain A into a domain B. In this work, domain A consists of 3D microscopy images of retinal vessels, and domain B corresponds to 3D segmentation masks of vessels. To the best of our knowledge, this is the first work proposing an automated approach for segmentation and quantification of 3D retinal blood vessels which does not require pairs of images and corresponding ground-truth masks for training. The source code of 3DVascNet is freely available, thus it can be easily extended to analyze other 3D vasculatures. Moreover, we provide a graphical user interface (GUI) suitable for researchers without a solid knowledge of programming.

The main contributions of this work are the following:

- We propose a DL-based approach for 3D retinal vessel segmentation in microscopy images that does not require paired 3D images and 3D masks for training;
- 3DVascNet, available as source-code and GUI, can be used for automated and accurate quantification of the 3D retinal vasculature;
- We demonstrate the robustness of our approach by testing it on four different datasets of mouse retinas;
- We release four datasets containing 3D microscopy images of retinal blood vessels, and corresponding 2D and 3D masks.

## 2 Methods

### 2.1 Datasets

The datasets used in this work were originally acquired and analyzed in a previous work [6]. The first dataset comprises 21 3D microscopy images of mouse retinal vessels, the corresponding 2D MIP images, and the corresponding 2D masks obtained by applying a classical segmentation pipeline to the MIP images as described in [18], which includes: thresholding, outliers and unconnected objects removal, morphological operations (binary erosion and dilation) and manual correction.

This dataset also contains 3D masks that were obtained from the 2D masks using a set of assumptions as proposed in [18]. Firstly, the skeletons and radii of each vessel segment were computed from the 2D masks. Retinal vessel skeleton is the representation of the vascular vessels as a one-pixel wide line containing pixels that are equidistant from the vessel boundaries, which is also denoted as the vessel centerline. Afterwards, the 3D masks were generated considering that the vessels’ 3D structure can be approximated as tubular segments, where each segment is cylindrical with a circular cross-section. The radius of each vessel segment in 3D was presumed to be equal to the radius computed from the corresponding 2D projection. Since these 3D masks were not obtained directly from the 3D images there is no overlap between these two across the z-axis. The only paired data are the 2D MIP images and the 2D masks. The 3D masks are a coarse representation of the vessels in the 3D images.

Table 1 describes dataset 1, where the prefix in each image name (first column) denotes a specific treatment provided to the retinas as described in the Table’s caption. We used a single image from each condition to train the 3D CycleGAN model, the remaining images were used as test data to evaluate the model’s performance.

**Table 1:** Description of dataset 1 used in this work for training and testing. The prefix in each image name (first column) represents the treatment that was applied to the retina. Retinas were collected from animals treated with: a) intraperitoneal injection of phosphate-buffered saline (PBS), which works as a control condition; b) intra-ocular injection of soluble fms-like tyrosine kinase 1 (sFLT1), a protein that blocks vascular endothelial growth factor A (VEGFA), which acts as an antiangiogenic molecule; c) intra-ocular injection of VEGFA, a pro-angiogenic molecule; and d) intraperitoneal injection of angiotensin-II (AngII), a vasoconstrictor peptide, which increases blood pressure. These images were previously collected and analysed in [6].

| Image | Size ($X \times Y \times Z$) | Train/Test Split |
| --- | --- | --- |
| PBS_ret1 | 5409×2979×31 | Train |
| PBS_ret2 | 3375×4599×56 | Test |
| PBS_ret3 | 6237×3384×38 | Test |
| PBS_ret4 | 5013×2979×43 | Test |
| PBS_ret5 | 4838×5561×39 | Test |
| PBS_ret6 | 4864×4864×88 | Test |
| PBS_ret7 | 6144×8192×20 | Test |
| sFLT1_ret1 | 4617×5418×35 | Train |
| sFLT1_ret2 | 4365×3978×73 | Test |
| sFLT1_ret3 | 4365×3600×66 | Test |
| VEGF_ret1 | 2979×4599×42 | Train |
| VEGF_ret2 | 5319×4878×61 | Test |
| VEGF_ret3 | 5427×4194×46 | Test |
| VEGF_ret4 | 5013×4194×38 | Test |
| VEGF_ret5 | 4365×3978×40 | Test |
| VEGF_ret6 | 4743×4365×61 | Test |
| VEGF_ret7 | 4365×4743×42 | Test |
| AngII_ret1 | 6273×3780×33 | Train |
| AngII_ret2 | 4851×5571×42 | Test |
| AngII_ret3 | 4864×3584×48 | Test |
| AngII_ret4 | 4864×5120×40 | Test |

Furthermore, to test the generalization capability of our segmentation model, we used another three datasets (datasets 2, 3 and 4) containing 28 3D images and 2D masks (Sup. Table 1). These datasets contain images of retinas that have undergone treatments other than those listed in Table 1, and were acquired by different users from those who obtained dataset 1. Additionally, 7 out of these 28 images were acquired in another laboratory, and consequently using a different microscope from the one used to obtain the training images.

All datasets will be made publicly available at https://huggingface.co/ datasets/Hemaxi/3DVesselSegmentation.

### 2.2 3DVascNet’s Pipeline

The proposed pipeline is depicted schematically in Fig. 1a. It consists of four main modules: 1) image pre-processing to enhance the vessels, 2) image segmentation based on the 3D CycleGAN model, 3) mask skeletonization, and 4) feature extraction based on the masks and skeletons. More information on individual blocks of the pipeline is provided in the following subsections.

**Fig. 1:**
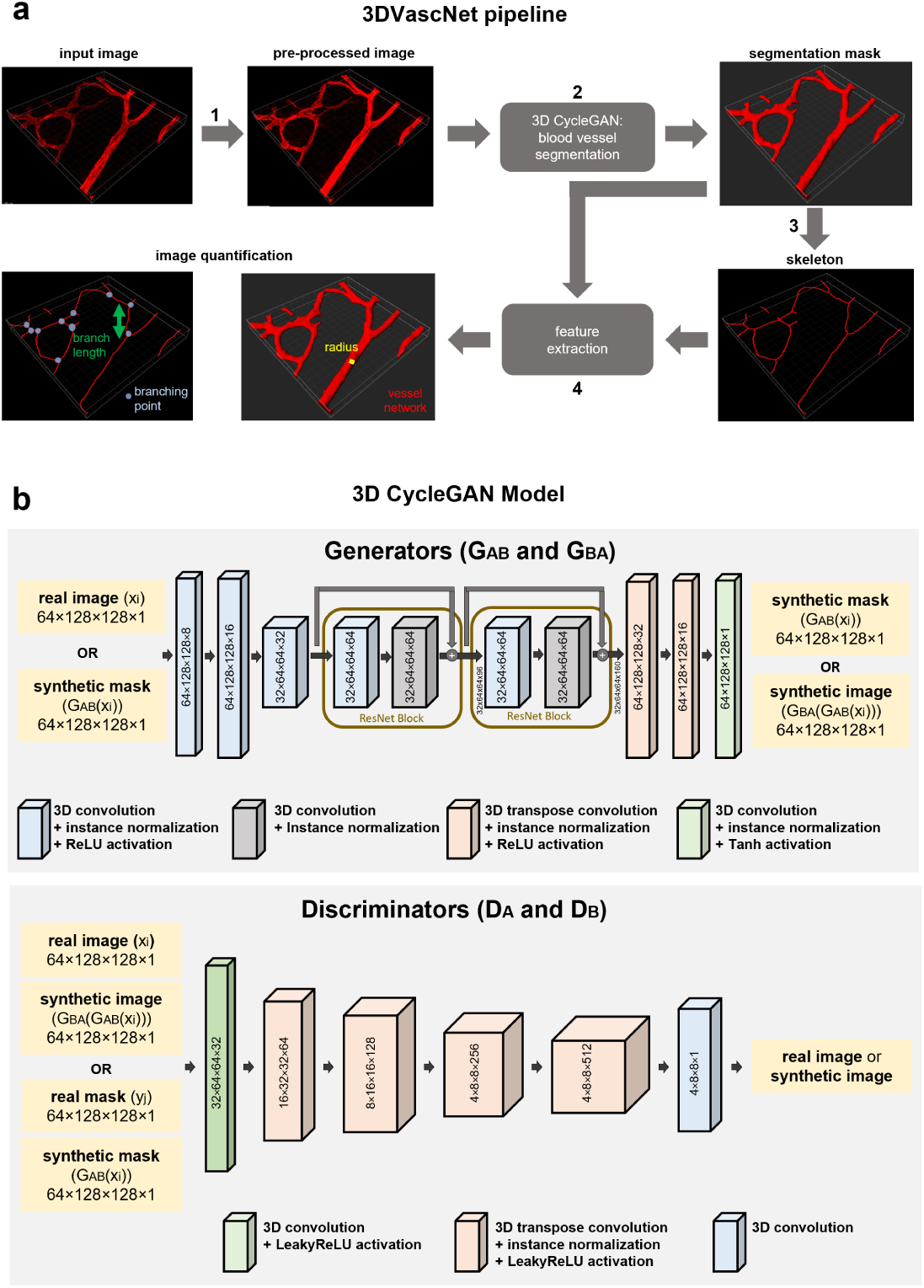
3DVascNet’s pipeline and 3D CycleGAN model architecture. **a** Overview of the 3DVascNet pipeline which is composed of four main blocks: 1) image pre-processing; 2) image segmentation based on the 3D Cycle-GAN model; 3) mask skeletonization; 4) feature extraction (vessel density, vessel radius, branch length, branching points density). **b** Architecture of the 3D CycleGAN model for retinal vessel segmentation in microscopy images. It is composed of two generators (top) and two discriminators (bottom). The numbers in the boxes denote the size of the feature maps resulting from the operation performed by the box.

#### 2.2.1 Pre-Processing

Several pre-processing methods have been proposed for intensity inhomogeneity correction, image normalization, and vessel enhancement to improve retinal vessel segmentation [11, 13, 19]. Here, similar to [8, 10], we performed normalization based on the 1 and 99 percentiles of the image intensity values. In this way, the voxels in the 1-99% range of intensity values are linearly mapped into the [0, 1] interval. Moreover, the bottom and top 1% of voxels are clipped at 0 and 1, respectively. Supplementary Fig. 1a shows an example of a patch extracted from the original microscopy image, and the corresponding patch extracted from the pre-processed image.

#### 2.2.2 3D Blood Vessel Segmentation

In this section, we present the proposed DL-based approach for 3D blood vessel segmentation, and its training and testing details. To overcome the problem of manually annotating 3D masks for training, in this work we propose a 3D CycleGAN model which only requires examples of images from the source domain and target domain for training. It does not require pairs of 3D images and masks. The source domain consists of 3D microscopy images, and the target domain comprises 3D masks that were created based on 2D masks of retinal vessels using the approach proposed in [18], as explained in subsection 2.1. Thus, manual annotation of 3D blood vessels is not needed in our approach.

##### 3D CycleGAN Model

The proposed 3D CycleGAN model is an extension to 3D of the 2D CycleGAN model developed for unpaired image-to-image translation [17]. To implement the 3D CycleGAN based on the 2D CycleGAN [17] we replaced the 2D layers (convolutional and transposed convolutional) by 3D layers. The 3D CycleGAN is composed of two generators (*{G_AB_, G_BA_}*), and two discriminators (*{D_A_, D_B_}*), where *G_AB_* maps volumes from domain A (source domain) into domain B (target domain), *G_BA_* translates volumes from domain B into domain A, and *D_A_*and *D_B_* classify, respectively, volumes from domains A and B as real or synthetic. The architecture of the generators and discriminators is depicted in Fig. 1b.

Each generator contains 3D convolutional, instance normalization and *ReLU* activation layers followed by a downsampling layer which reduces the size of the feature maps. Thereafter, it contains residual blocks which are composed of 3D convolutions, instance normalization and *ReLU* activation. Finally, an upsampling layer increases the size of the feature maps, and it is followed by 3D transposed convolution, instance normalization and *ReLU* activation layers, except the activation function of the last layer which is a *Tanh* activation function as proposed in [17]. The discriminator is a convolutional neural network designed for image classification. It takes as input a real or synthetic volume and outputs the probability of this volume being real or synthetic. The discriminator is composed of 3D convolutional, instance normalization and *LeakyReLU* activation layers (Fig. 1b).

Considering an unpaired dataset containing volumes of microscopy images 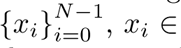 domain A) and volumes of masks (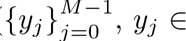 domain B). The mapping *G_AB_*: *A → B* receives an image *x_i_ ∈* domain A and learns to generate a mask *G_AB_*(*x_i_*) that is indistinguishable from masks *y_j_ ∈* domain B. It learns this mapping by adversarial training as proposed in [20]. That is, on the one hand, *G_AB_* learns to generate synthetic masks, on the other hand *D_B_* tries to distinguish real masks (*y_j_*) from synthetic ones (*G_AB_*(*x_i_*)), and the goal of *G_AB_* is to fool the discriminator *D_B_* by generating synthetic masks similar to the real ones. Nevertheless, this does not ensure that *x_i_* and *G_AB_*(*x_i_*) will be paired. Therefore, structure is added to the 3D CycleGAN to encourage that if we transform an image (*x_i_*) into a mask (*G_AB_*(*x_i_*)) and then transform it back into an image (*G_BA_*(*G_AB_*(*x_i_*)) we get an image similar to *x_i_*. This is expressed as: *G_BA_*(*G_AB_*(*x_i_*)) *≈ x_i_*, where *G_BA_*: *B → A* is the mapping from the masks set into the images set. To add this structure, the 3D CycleGAN has two training paths: forward (*x_i_→ G_AB_*(*x_i_*) *→ G_BA_*(*G_AB_*(*x_i_*)) *≈ x_i_*) and backward (*y_j_ → G_BA_*(*y_j_*) *→ G_AB_*(*G_BA_*(*y_j_*)) *≈ y_j_*), as represented in Sup. Fig. 1b top).

Thus, mappings *G_AB_*: *A → B* and *G_BA_*: *B → A* along with their respective generators, *D_B_* and *D_A_*, are trained through adversarial training. More specifically, the training process is cyclical where, at each iteration, *G_BA_*: *B → A* is updated, followed by training *D_A_* using the generated samples, then *G_AB_*: *A → B* is updated, followed by updating *D_B_* using the generated samples. Moreover, the addition of the cycle consistency loss encourages *G_AB_*(*G_BA_*(*y_j_*)) *≈ y_j_* and *G_BA_*(*G_AB_*(*x_i_*)) *≈ x_i_*. Finally, an identity term is added to preserve the color. Hence, the objective function of the 3D CycleGAN contains four terms:

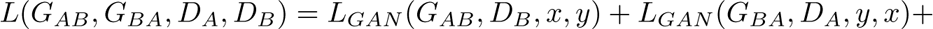

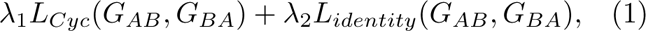

where *λ*_1_ and *λ*_2_ are constants. The first two terms are the adversarial losses:

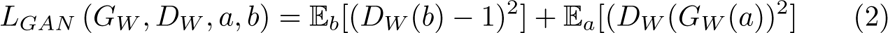

where E*_a_*and E*_b_* represent the expectation of the log-likelihood of a sample being sampled from the probability distribution of *a* and *b*, respectively. The goal of *G_W_* is to maximize this objective and the goal of *D_W_* to minimize it: max*_GW_* min*_DW_ L_GAN_* (*G_W_, D_W_, a, b*). In this work, the loss shown in Eq. 2 is applied to the pairs *{G_AB_, D_B_}* and *{G_BA_, D_A_}*.

The third term is the cycle consistency loss:

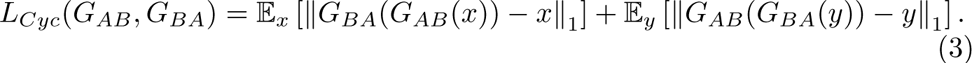

The last term is the identity loss:

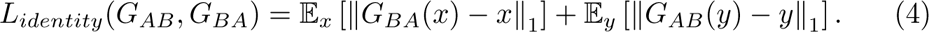

Our implementation of the 3D CycleGAN is made publicly available at https://github.com/HemaxiN/3DVascNet, it is based on a 2D implementation by J. Brownlee ^1^.

##### Training

To train the 3D CycleGAN model we extracted non-overlapping volumes of size 128 *×* 128 *×* 64 from the 3D images and 3D masks. Padding was performed before extracting the volumes, and the new dimensions 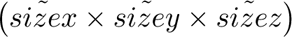 of the images/masks were computed as follows:

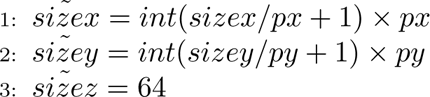

where *sizex* and *sizey* denote the original sizes of the image/mask along the x and y dimensions, *int* (*·*) converts the specified value into the largest integer not greater than the value, *px* and *py* denote the patch size along the *x* and *y* axes, respectively. In this way, the image size is adapted to be a multiple of the patch size along x and y. For the 3D masks we performed zero padding. The original and pre-processed 3D images were padded, respectively, with small random values in the interval [1, 25] and [1, 50], since these are the typical voxel intensity values in the background of these images.

After performing padding and extracting volumes from the images and masks, we obtained 4987 image and 4987 mask volumes for training, which were normalized to present values ranging from -1 to 1. Based on our experiments, we concluded that training the model for a large number of epochs deteriorated the results. For instance, after 11 epochs the output of the model was a tensor of zeros (Sup. Fig. 2a). The best models were obtained between epochs 1 and 5. We trained the 3D CycleGAN from scratch for 15 epochs, using Adam optimizer (learning rate of 1e-4 and *β*_1_ = 0.5) [21], and setting *λ*_1_ = 10 and *λ*_2_ = 5 as suggested in [17] and defined in Eq. 1. The batch size was set to 1 due to memory limitations. Training the 3D CycleGAN with this configuration took approximately 36 hours. The best epoch was selected based on visual inspection of the segmentation masks at each epoch, and considering an unsupervised evaluation metric that is described in subsection 2.4 (Sup. Fig. 2).

##### Testing

For testing we extracted volumes of size 128 *×* 128 *×* 64 with an overlap of 64 voxels along the *x* and *y* directions, and 32 voxels along the *z*-direction. These volumes were extracted from the padded images. For the test images, padding was performed as explained above, except for images with more than 64 slices. For these images the new dimension along z was computed as follows: 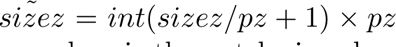, where *sizez* is the original image size across z, and *pz* is the patch size along the z-axis. The volumes were then fed to the trained generator *G_AB_*: *A → B* (Sup. Fig. 1b bottom) which outputs volumes (*out*) with values ranging from -1 to 1. Firstly, the following operation was performed to ensure that the output volume values are within the range [0, 1]: 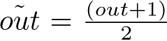. Afterwards, a threshold of 0.5 was applied to *oũt*: voxels with values less than 0.5 were set to zero, indicating background, while all others were set to one, representing the vessels. In order to incorporate patches with overlapping regions, we employed a voting mechanism based on two variables: *N_back_* and *N_fore_*. Here, *N_back_*[*u, v, x*] and *N_fore_*[*u, v, x*] count the occurrences of voxel (*u, v, x*) being classified as background and foreground, respectively. Thereafter, the voxel values in the output segmentation mask were computed as follows:

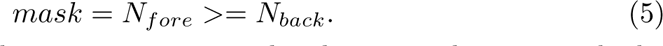

Finally, we cropped the segmentation masks along x and y to match the size of the corresponding images. The test time for about 37 thousand patches of size (64, 128, 128) is approximately 1 hour.

##### Post-Processing

After performing vessels segmentation, we applied a simple post-processing step. It is known that the vessels in the retina are connected, thus they must be represented as a binary tree-like configuration in the segmentation mask [12]. Hence, as a post-processing step we removed unconnected objects in the output segmentation mask, thus the final segmentation mask contains only the largest connected object.

#### 2.2.3 Definition of a Region of Interest (ROI)

The segmentation masks typically contain the vascular structure and background regions. Thus, we need to define a ROI to extract features only from the vascular network. This allows for more targeted analysis and avoids unnecessary computations on irrelevant regions.

In this work, we automatically defined the 3D ROIs for each image. Firstly, we obtained the 3D convex hull from each 3D segmentation mask. Thereafter, we computed the concave hull of each 2D segmentation mask [22], and applied some pre- and post-processing steps (Sup. Fig. 3a). The 2D masks were obtained from the 3D segmentation masks by performing an AND operation along the z-direction. To compute the concave hull, firstly the segmentation mask was resized to (100,100) to decrease the computational cost. Then, the coordinates of the points belonging to the vessel edges were used to compute the concave hull as proposed in [22]. Thereafter, the resulting binary mask was resized to the original size, and two post-processing steps were applied. Firstly, regions outside the concave hull that were smaller than 5% of the largest region outside the concave hull were added to the ROI. Thereafter, median filter was applied to reduce staircase-like borders shown in Sup. Fig. 3a. Lastly, to create the final 3D ROI, we performed an intersection of the 2D ROI with each slice along the z-direction obtained from the 3D convex hull (Sup. Video 1).

Vascular biologists typically perform selective feature extraction to focus on specific areas of interest. Firstly, they define several ROIs on the images, thereafter they extract features from these ROIs to study vascular networks. While our automated approach computes a single ROI for the input image, we have an optional step in our GUI that allows users to manually select multiple ROIs.

#### 2.2.4 Skeletonization

After obtaining the vessels’ segmentation masks and ROIs, the next step consists in extracting the vessels’ skeletons. Before computing the skeleton, we applied a 3D morphological closing operation with a binary ellipsoid of size (9,9,3) to fill small holes in the 3D segmentation masks.

A 3D segmentation mask and the corresponding retinal vessel skeleton are represented in Sup. Fig. 3b. A branching point is a point connected to three or more vessel segments, an endpoint is a point connected to one vessel segment (Sup. Fig. 3b). A branch is a segment between two branching points, or between an endpoint and a branching point.

In this study we only have 2D ground-truth masks, thus we extracted 2D skeletons from the predicted and ground-truth masks to compare retinal vasculature quantification. The predicted 3D segmentation masks were projected into 2D using an AND operation along the z-axis. To avoid false positive centerlines, as proposed in [10], we applied erosion followed by dilation to the 2D masks using an ellipsoid structuring element of size (5, 5). Then, the skeleton of each segmentation mask was computed inside the ROI using the method proposed by Lee et al. [23].

However, as explained in section 1, 2D skeletons do not accurately represent the real structure of 3D vessels. Herein, they were computed only for comparison purposes. Our 3DVascNet software computes the 3D skeletons from the 3D segmentation masks using the method presented in [23], to perform retinal vasculature quantification in 3D.

#### 2.2.5 Retinal Vasculature Features

In this work, based on both the segmentation masks and their skeletons, we extracted four features to describe retinal blood vessels: vessel density (vascular volume/ROI volume), mean vessel radius, mean branch length and branching points density.

For each image, the four above-mentioned features were computed within the ROI:

- **Vessel Density (Vascular Area/ROI Area)** - We compute the quantity 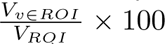, where *V_ROI_* and *V_v∈ROI_* denote the volume (number of voxels) of the ROI and of the vessels inside the ROI, respectively.
- **Mean Vessel Radius** - Briefly, for each branch in the skeleton, its direction was computed, and the vessel cross-section orthogonal to that direction was determined for each point (*p*) in the branch. The vessel contour was extracted from the cross-section, which is composed of N points (*c*_0_*, c*_1_*, …, c_N−_*_1_). The mean vessel radius for each point in the branch was computed as the mean of the Euclidean distances between each contour point and the branch point: 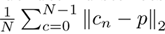. The mean vessel radius corresponds to the mean of the radii computed for all points in all branches.
- **Branching Points Density** - We compute 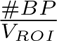, where #*BP* denotes the number of branching points.
- **Mean Branch Length** - It represents the mean of the individual branches’ lengths. That is, it is the mean of the sum of the Euclidean distances between consecutive voxels along each branch in the skeleton.

The length of the branches and the number of bifurcation points were computed using the Skan tool [24]. The voxel spacing was taken into consideration to compute the branch lengths and vessel radii in *µm*. These features were also computed in 2D for comparison purposes.

### 2.3 Compared Approaches

The approaches that we compared with are the following:

- Otsu: It is an easy and fast method to segment images. It selects a threshold based on the histogram of image intensities. The threshold is selected in order to maximize inter-class variance [13, 25, 26]. For a bimodal histogram the selected threshold separates the two histogram peaks.
- Chan-Vese [27]: It is an iterative algorithm that belongs to a family of methods for image segmentation named geometric active contours or snakes. Chan-Vese works with the assumption that the foreground and background pixel intensity values follow two Gaussian distributions. Thus, it presents good results when the background and foreground regions have different means. Segmentation is obtained by minimizing an energy function, which is the weighted sum of four terms (Eq. 6): (i) the sum of the squared differences between the intensities of the voxels in the segmented region and the mean intensity of this region, (ii) the sum of the squared differences between the intensities of the voxels in the background region and the mean intensity of this region, (iii) a term dependent on the volume of the segmented region and (iv) another term dependent on the length of the segmented region’s boundary:

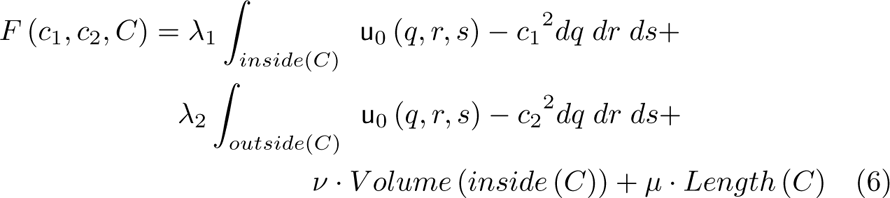

where *u*_0_ denotes the image to be segmented, *C* the curve that encloses the foreground region, *c*_1_ and *c*_2_ are the averages of the voxel intensity values inside and outside *C*, respectively; and *ν ≥* 0, *µ ≥* 0, *λ*_1_ *>* 0 and *λ*_2_ *>* 0 are the weights applied to each term.

To test this algorithm in our dataset we used a publicly available implementation: https://github.com/pmneila/morphsnakes.

### 2.4 Evaluation Metrics

Several supervised evaluation metrics have been proposed for the quantitative assessment of retinal vessel segmentation approaches: Dice coefficient (DC) [28], accuracy, sensitivity, and specificity, among others. In these metrics, the segmentation masks obtained with a given method are compared with the respective ground-truth masks which are typically manually annotated by experts [11–13, 19, 29]. Herein, we used the DC metric [28]:

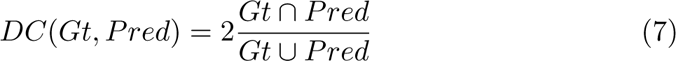

which is computed as a ratio of the intersection between the ground-truth (Gt)and predicted masks (Pred) by the union of Gt and Pred masks. However, this supervised metric requires manual segmentation masks which are difficult to obtain. Besides, it may be subjective due to the inter-expert variability associated to manual segmentation [30]. Herein, since we already had the 2D masks, that were generated in a previous study [6], we used those to compute the DC values to evaluate the performance of the compared segmentation approaches.

Unsupervised evaluation metrics do not require ground-truth segmentation masks. Instead, these metrics are computed based on the comparison between each segmentation mask generated by a given model and the corresponding original input image. Zhang et al. [30] evaluated the performance of several unsupervised metrics, such as entropy, texture and color error. They computed the accuracy for each unsupervised metric, which represents the number of times the metric gives a better score to the mask annotated by a human evaluator instead of the mask generated by a segmentation model [30].

Unsupervised metrics can be computed for any segmented image and are objective [30]. Nevertheless, these metrics have disadvantages, for instance, given two masks A (optimal segmentation) and B (noisy segmentation) an unsupervised metric does not always give a better segmentation score to mask Zhang et al. [30] obtained accuracy values between 1.0% and 82.1% for the studied unsupervised evaluation metrics.

In this work, to select the best epoch of the 3D CycleGAN model, we propose the normalized mutual information (NMI) [31] as an unsupervised segmentation evaluation metric, since it is easier to compute than the metrics studied in [30]. NMI is a widely used metric to measure the alignment between two images of different modalities, without requiring the signals to be identical in these two images. It measures the predictability of the signal in one image given the signal in another image. This metric is appropriate to measure the level of similarity between the original microscopy images, with values in the interval [0, 255], and the corresponding binary segmentation masks with zeros and ones, because it assesses how well the segmentation captures the vascular structures present in the image (Sup. Fig. 2). NMI is computed as follows:

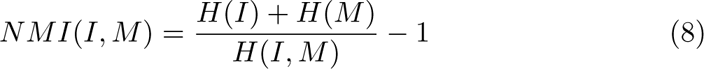

where:

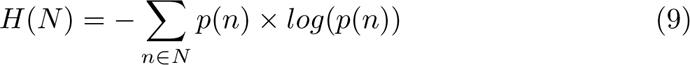

is the marginal entropy, it represents the average information for an image (*N*) containing the set of values *n* (*n ∈ N*), and *p*(*n*) denotes the marginal probabilities of individual values [31], and:

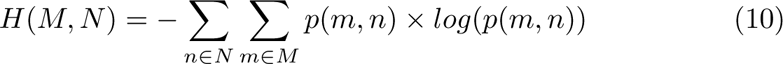

is the joint entropy of images *M* and *N*, and the joint probability *p*(*m, n*) determines how frequently pairs of values occur together [31].

### 2.5 Statistical Analysis

The data were tested with a Mann-Whitney U test, p-value annotation legend: ‘ns’ 0.05 *< p − value <*= 1, * 0.01 *< p − value <*= 0.05, ** 0.001 *< p − value <*= 0.01, *** 0.0001 *< p − value <*= 0.001. Correlation was studied using linear regression. All statistical analyses were performed in Python.

### 2.6 Computer Specifications

The experiments were performed in Python 3.10 on a computer equipped with an NVIDIA GPU GeForce 1080 Ti. The 3D CycleGAN was trained using the open-source DL library Keras [32]. We run all experiments using the sourcecode, and we provide a GUI for researchers without a solid programming background.

### 2.7 3DVascNet’s GUI

A detailed section explaining the installation and step-by-step use of 3DVascNet’s GUI can be found in Supplementary Material I. Briefly, the directory containing the 3D images must be selected by clicking on the “Load the Image(s)” button (Sup. Fig. 4a and b). These images will be segmented and afterwards quantification will be performed upon clicking on the “Segment” and then “Quantify” buttons, respectively. During this analysis a results folder will be automatically created in the images folder (Sup. Fig. 4c). The segmentation masks and skeletons of the images will be saved in the results folder. These can be visualized using the horizontal and vertical sliders (Sup. Fig. 4d). Although our software automatically computes the ROI, there is an option to manually select one or more ROIs (”Select ROI” button) (Sup. Fig. 4e). After processing all images, a CSV file containing all morphological features for each image will also be saved in the results folder upon clicking on the “Save the Results” button (Sup. Fig. 4f). Afterwards, the distributions of the features for different groups can be visualized by specifying the group names (separated by commas and without spaces, for instance: Captopril,VEGF,PBS), by selecting the features to be visualized and using the “Visualize Boxplots” button (Sup. Fig. 4g). To load a previously generated CSV file, the “Load CSV” button can be used at any time, and vasculature features’ distributions can be visualized without requiring to perform segmentation and quantification. Finally, the “3D Visualization” button allows visualization in 3D of the images and/or segmentation masks based on the napari viewer [33].

## 3 Results

### 3.1 DL-Based Segmentation Model

Most DL-based approaches require large, manually annotated, ground-truth datasets for training and validation. These annotations should be sufficiently accurate for the model to perform correctly. However, manual segmentation of blood vessels in 3D is very demanding. The generation of 3D datasets of sufficient diversity and complexity would require a large effort from researchers. Thus, to overcome the necessity of manually annotating 3D segmentation masks for training, we explored a specific DL method that only requires examples of images from the source domain and target domain for training, the so-called 3D CycleGAN model [17]. We trained the 3D CycleGAN model with 3D microscopy images of retinal blood vessels (the source domain), and synthetic 3D masks (the target domain). The synthetic 3D masks were created based on PolNet [18], using manual 2D masks of retinal vessels, which were previously generated and published by Barbacena et al. [6]. This way we overcome the burden and need of generating manually annotated 3D masks.

To select the best epoch of the 3D CycleGAN model we used the NMI metric [31]. Inspired by the work of Zhang et al. [30], we evaluated the performance of the NMI metric computed between 2D images of blood vessels and 2D masks (ground-truth masks and masks with some perturbations). For each ground-truth segmentation mask, we generated four masks by applying: dilation and erosion (considering an ellipsoid structuring element of size (5, 5)), and salt and pepper noise with probability 0.1 and 0.04. Thereafter, for each image, we computed 5 values of NMI (Eq. 8) between the image and each of the five masks (the ground-truth mask and the four transformed masks). Finally, we computed the accuracy of the NMI metric as the number of times this metric gives a better score (higher value) to the ideal mask (annotated by experts) instead of the transformed mask. Our results showed that NMI is a good unsupervised evaluation metric since it presents high accuracy values which means that it performs well in giving a better score to the best segmentation mask (Table 2). Remarkably, the accuracy values obtained for NMI in our dataset are considerably higher than the ones obtained with the metrics and datasets considered in [30].

**Table 2:** Accuracy (%) of the unsupervised segmentation evaluation metric: NMI (Eq. 8).

| 2D Mask with Perturbations | Dilation | Erosion | Salt and Pepper Noise (p=0.1) | Salt and Pepper Noise (p=0.04) |
| --- | --- | --- | --- | --- |
| Original Image | 93.88% | 85.71% | 100% | 100% |
| Normalized Image | 93.88% | 100% | 100% | 100% |

Next, we studied the performance of the 3D CycleGAN model using preprocessing and post-processing steps. Figure 2a shows the segmentation masks obtained with the 3D CycleGAN model for a raw and a pre-processed image patch (two masks on the top), and the corresponding post-processed masks (two masks on the bottom). These results show that the post-processing step improves the quality of both segmentation masks by removing noise and maintaining only the largest connected object. However, even after applying the post-processing step, the segmentation mask obtained with the raw image presents some errors, such as oversegmentation of vessels, indicated with a white arrow. Moreover, our quantitative results on the test images from dataset 1 showed that the 3D CycleGAN model tested on the raw and pre-processed images, with the post-processing step, achieves DC values of 0.8720*±*0.0367 and 0.9319*±*0.0288, respectively. Hence, the qualitative and quantitative results suggest that the best segmentation masks are obtained considering both the pre- and post-processing steps. These results show that we generated a DL-based approach that is capable of accurately segmenting blood vessels in 3D without needing 3D manual ground-truth for training.

**Fig. 2:**
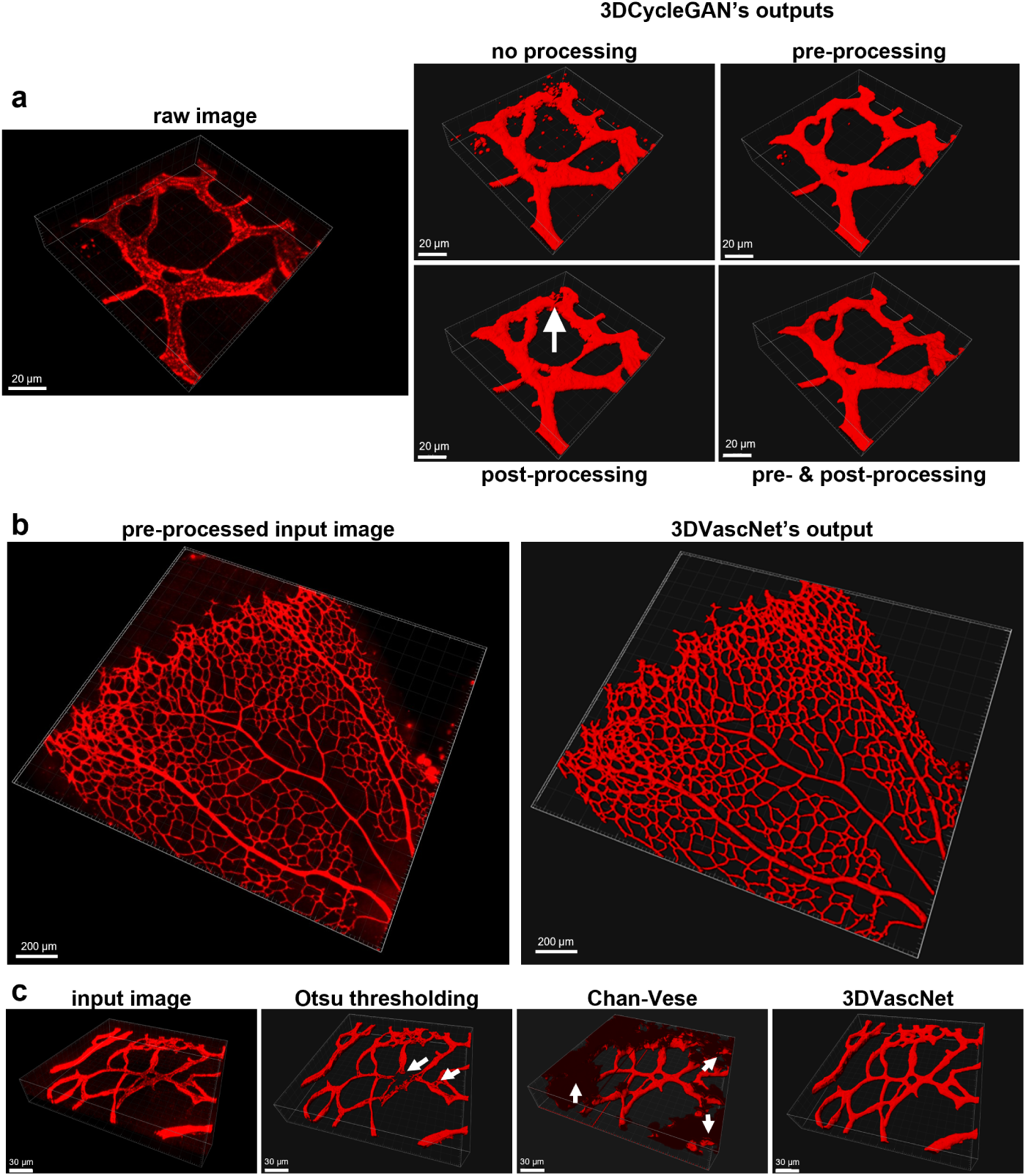
Optimization of the 3DVascNet model and comparative performance. **a** Demonstration of the effects of pre-processing and postprocessing steps in the output of 3DVascNet using as an example a 3D patch extracted from the input image (left), and the corresponding segmentation masks obtained with the 3DVascNet (right), corresponding to the different processing steps. **b** Example of the segmentation of an entire P6 retinal vascular network, labelled by ICAM2, using 3DVascNet. Input image (left), and 3DVascNet segmentation mask (right). **c** Examples of a 3D patch extracted from an input image (left), and the corresponding segmentation masks obtained with Otsu thresholding, Chan-Vese and 3DVascNet.

We named our approach 3DVascNet. A full-scale image of the segmentation obtained using 3DVascNet is shown in Fig. 2b.

Next, we evaluated our method and pipeline by comparing the performance of 3DVascNet with two classical methods for image segmentation. Since our dataset does not contain 3D ground-truth masks, we are unable to compare the performance of 3DVascNet with supervised segmentation methods. Thus, we considered two unsupervised methods for segmentation: Otsu thresholding [13, 25, 26] and Chan-Vese [27]. These approaches were applied on the pre-processed images, and the output segmentation masks were subjected to the post-processing step. For qualitative comparison we show in Fig. 2c an example of a patch extracted from a 3D pre-processed image, and corresponding segmentation masks obtained with Otsu thresholding, Chan-Vese and 3DVascNet. On the one hand, Otsu thresholding is more sensitive to heterogeneous staining resulting in errors in the segmentation mask as shown with white arrows (Fig. 2c). On the other hand, Chan-Vese is more susceptible to the background noise compared to Otsu thresholding and 3DVascNet (white arrows in Fig. 2c). More examples of segmentation masks obtained with the compared approaches are shown in Sup. Fig. 5. The qualitative results indicate that the best masks are obtained with 3DVascNet (Fig. 2c and Sup. Fig. 5). The quantitative results, represented in Table 4, are consistent with the qualitative results. In fact, 3DVascNet achieves the highest DC values (Table 4): DC values of 0.9450 ± 0.0240 and 0.9319 ± 0.0288, on the training and test sets, respectively.

### 3.2 Generalization Capability

We next evaluated the generalization capability of 3DVascNet. First, we measured the performance 3DVascNet on additional datasets of retinal vascular networks, acquired with different vessel markers, microscopes and by different users (datasets 2, 3 and 4 described in Sup. Table 1). Figure 3a represents the segmentation masks obtained with 3DVascNet for three patches taken from images from datasets 2-4. These results show that 3DVascNet provides segmentation masks with high-quality for images acquired under different conditions and on different microscopes. This is confirmed by comparing 3DVascNet with Otsu and Chan-Vese methods (Table 3). In fact, 3DVascNet outperforms the traditional methods for segmentation, presenting the highest DC mean and smallest standard deviation for all datasets considered. The 3DVascNet model achieves mean DC values of *≈*93% for these three datasets, whilst Otsu thresholding achieves mean DC values of *≈*88%, and Chan-Vese only *≈*76%. Hence, 3DVascNet presents a high generalization capability.

**Fig. 3:**
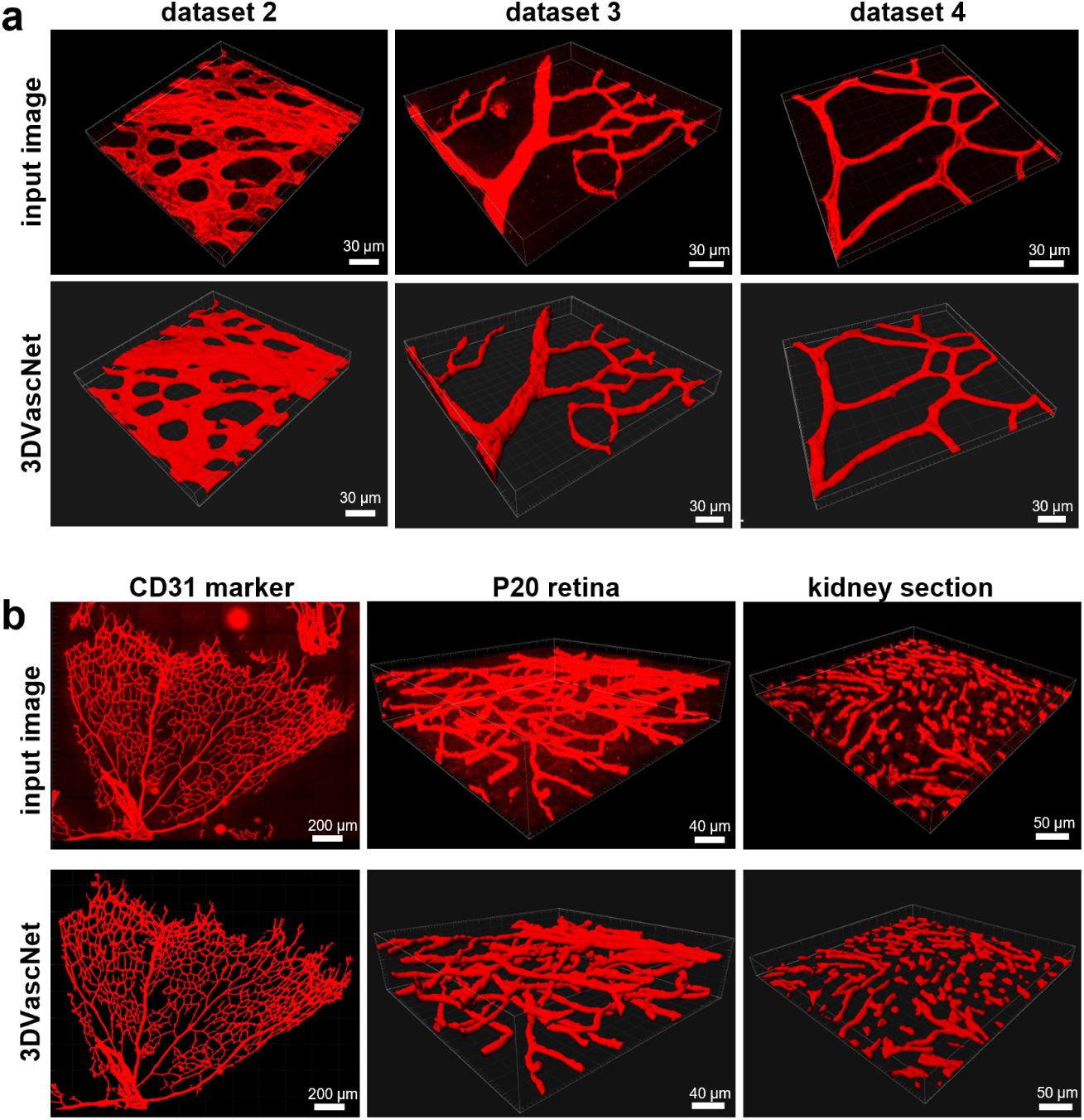
Generalization Capability of 3DVascNet. **a** Examples of 3D patches extracted from the input images (top), and the corresponding segmentation masks obtained with 3DVascNet (bottom), corresponding to datasets 2, 3 and 4 (from left to right, respectively). **b** Examples of 3D patches extracted from the input images (top), and the corresponding segmentation masks obtained with 3DVascNet (bottom), corresponding to different conditions (CD31 marker and P20 retina), and a different organ (kidney section).

**Table 3:** DC values of the compared vessel segmentation approaches on datasets 2, 3 and 4, considering the images after applying the percentile based normalization. All values correspond to *mean ± std*. The best results are high-lighted in **bold**.

| Test | Otsu | Chan-Vese | 3DVascNet |
| --- | --- | --- | --- |
| Dataset 2 | 0.9236 ± 0.0287 | 0.7947 ± 0.1271 | **0.9486 ± 0.0186** |
| Dataset 3 | 0.8228 ± 0.1545 | 0.6617 ± 0.2494 | **0.9230 ± 0.0816** |
| Dataset 4 | 0.9020 ± 0.0180 | 0.8743 ± 0.0550 | **0.9380 ± 0.0109** |
| All | 0.8750 ± 0.1106 | 0.7576 ± 0.1972 | **0.9349 ± 0.0545** |

**Table 4:**
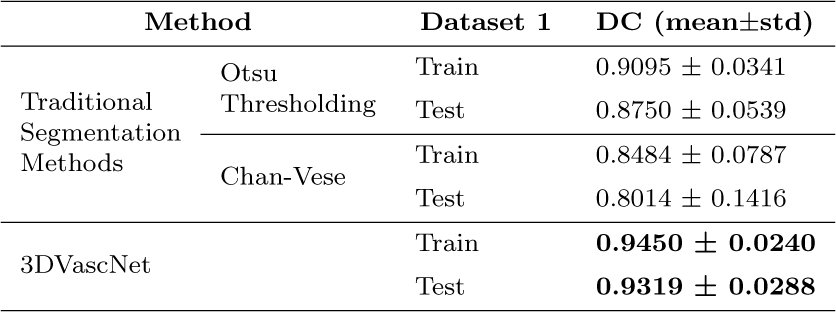
DC values of the compared vessel segmentation approaches on dataset 1, considering the pre-processing and post-processing steps. All values correspond to *mean ± std*. The best results are highlighted in **bold**.

Next, we qualitatively studied the generalization capability of the proposed segmentation model using additional experimental conditions. First, we tested if 3DVascNet could perform well using CD31 staining instead of ICAM2, which was used for the previous datasets (1-4). Indeed, CD31 staining was also well-segmented as ICAM2, both at the same stage of retinal development (postnatal day 6) as the previous datasets (Fig. 3b). 3DVascNet also segments well three layers of blood vessels corresponding to a later stage (post-natal day 20), where the three-dimensionality of the network is more apparent (Fig. 3b, Sup. Videos 2-4). In addition, we segmented a thick section from kidney medulla stained for CD31. Remarkably, 3DVascNet also performed very robustly and correctly segmented almost all individual vessels.

Thus, 3DVascNet presents a good ability to segment vessels, even in the presence of image variations resulting from differences in staining techniques, microscopy equipment, image modality, mouse organ and its developmental stage. Training with further datasets will likely further expand its generalization capabilities.

To facilitate the usage of 3DVascNet, we have generated a GUI that has been developed in Python (Sup. Fig. 4). 3DVascNet’s GUI allows automatic analysis of 3D microscopy images of blood vessels. 3DVascNet has been trained only with images from retinal blood vessels at post-natal day 6 of development. Thus, it should perform well on similar datasets. Yet, other vascular networks can be tested using our software.

### 3.3 Benchmarking 3DVascNet for Vascular Morphometrics

Next, we tested if the quantification of several vascular parameters (vascular morphometrics), such as vessel density, vessel radius, number of branching points and vessel length, using 3DVascNet were comparable with vascular morphometrics obtained from manual segmentation. To be able to have similar dimensions and units, we calculated vascular morphometrics from 2D MIP of the 3DVascNet’s masks and compared them with the vascular morphometrics computed from the 2D ground-truth masks. Remarkably, the distributions for the different vascular morphometrics obtained through 3DVascNet are very similar to the distributions obtained using user-dependent 2D groundtruth (Fig. 4). Importantly, there are no statistical differences, which confirms that results are equivalent between both approaches. To further confirm that results are within the same range, we also performed a Pearson correlation analysis between the values of vascular morphometrics based on the manual ground-truth masks and those based on the 3DVascNet’s masks. We obtained high *R*^2^ values for all features (*R*^2^ close to 0.9), which demonstrates a linear relationship between two variables.

**Fig. 4:**
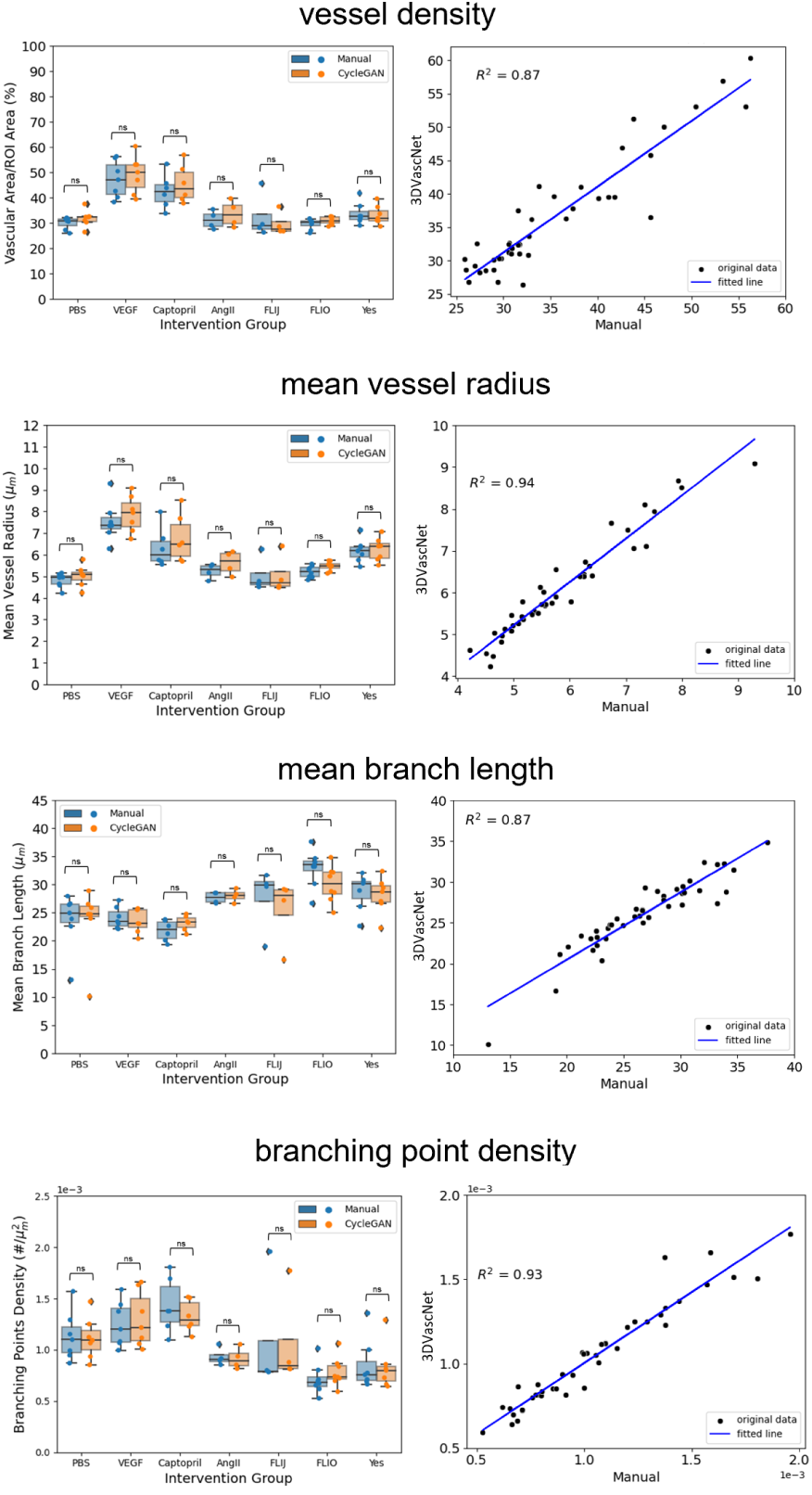
Performance of 3DVascNet using Vascular Morphometrics. Boxplots (left graph) represent side-by-side comparison of vascular morphometrics computed from 2D MIP of manual segmentation masks (in blue) or 3DVascNet masks (in orange). Corresponding Pearson correlation analysis (right graph) between the vascular morphometrics based on manual masks (x-axis) and those based on the 3DVascNet masks (y-axis). Data represents vessel density (vascular area/ROI area (%)); mean vessel radius (*µm*); mean branch length (*µm*) and branching point density (#*/µm*^2^). For correlation analyses, we considered 43 images, corresponding to the groups shown in the boxplots. Here we considered only the intervention groups that have at least 4 images: PBS (N=7), VEGF (N=7), Captopril (N=6), AngII (N=4), FLIJ (N=4), FLIO (N=8), Yes (N=7)). Statistical results from a Mann-Whitney U test. Statistical analysis legend: ‘ns’ 0.05 *< p − value <*= 1, * 0.01 *< p − value <*= 0.05, ** 0.001 *< p − value <*= 0.01, *** 0.0001 *< p − value <*= 0.001.

These findings demonstrate that 3DVascNet is a powerful automated method for blood vessels segmentation and quantification yielding robust scientific outcomes that are within the range of manual segmentation.

### 3.4 3D Quantification of Vascular Morphometrics

Next, we tested if we could obtain additional information from our datasets when using quantifications in 3D instead of quantifications from 2D MIP images. To do so, we computed 3D skeletons using the method proposed in [23] (Fig. 5a). We used the 3D skeletons to calculate the four previously characterized vascular morphometric parameters in 3D (Fig. 5b). Interestingly, our results demonstrate that some quantifications, and some relative differences between experimental groups, differ when quantifying parameters in 2D or 3D, whilst in other cases tendencies are maintained (Fig. 5b). For instance, the vessel density in 3D follows the same trends as 2D quantifications. However, we must note that absolute values differ very much, and 2D quantifications tend to suggest a much higher vessel density than in reality.

**Fig. 5:**
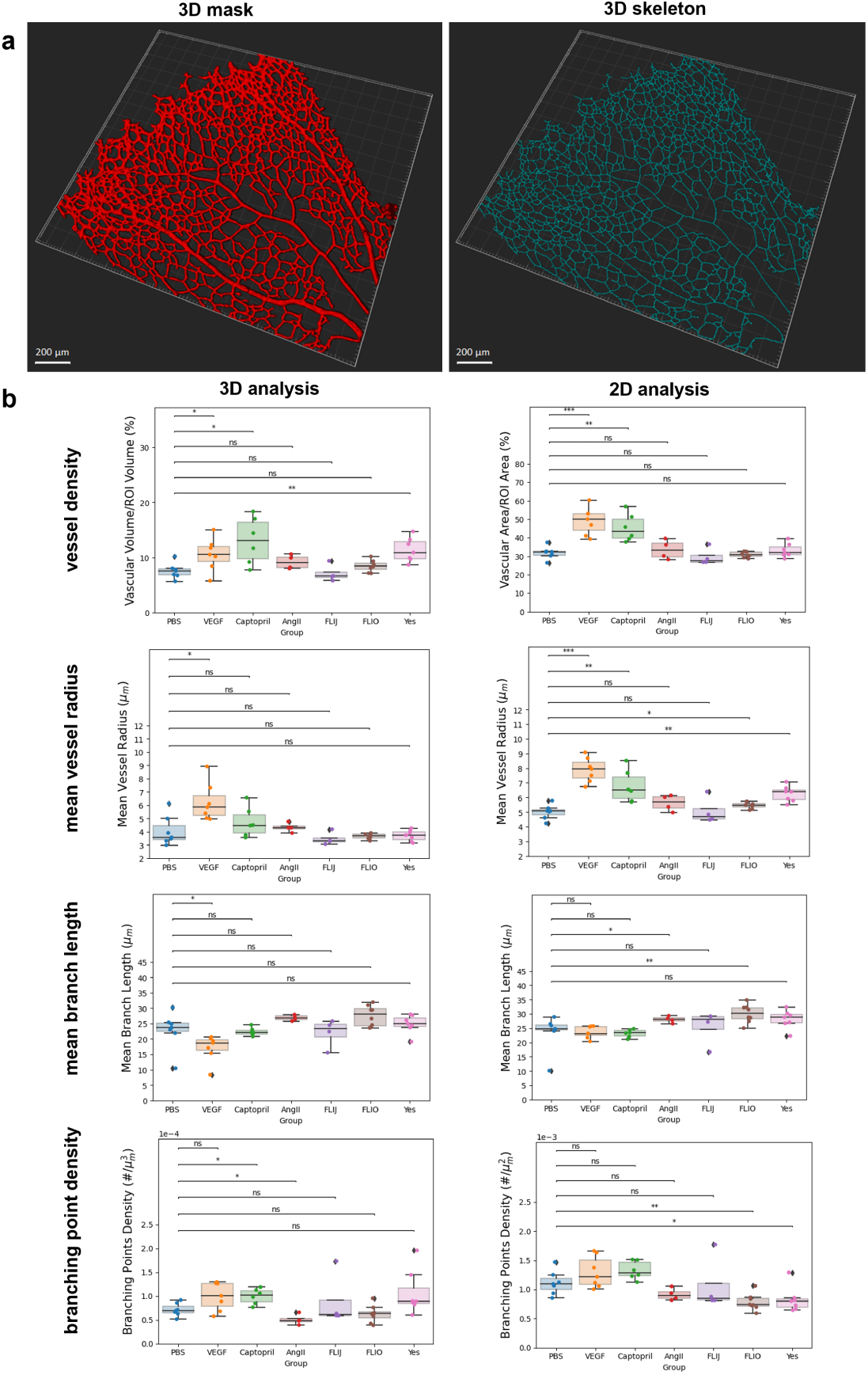
Comparison between 2D and 3D Vascular Morphometrics. **a** Representative 3D segmentation mask from an AngII retina (left image) and corresponding 3D skeleton (right image) computed based on the segmentation mask. **b** Boxplots represent vascular morphometrics computed from 3D VascNet’s masks (left graph) or 2D MIP of 3DVascNet’s masks (right graph), corresponding to the four studied morphometrics: vessel density (%); mean vessel radius (*µm*); mean branch length (*µm*) and branching point density (#*/µm*^3^ or #*/µm*^2^). Here we also considered only the intervention groups that have at least 4 images: PBS (N=7), VEGF (N=7), Captopril (N=6), AngII (N=4), FLIJ (N=4), FLIO (N=8), Yes (N=7)). Statistical results from a Mann-Whitney U test. Statistical analysis legend: ‘ns’ 0.05 *< p − value <*= 1, * 0.01 *< p − value <*= 0.05, ** 0.001 *< p − value <*= 0.01, *** 0.0001 *< p − value <*= 0.001.

Importantly, the mean vessel radius was substantially changed. Notably, higher mean vessel radius values were observed in 2D compared to 3D. Also, in 3D, most of the conditions do not present a significant difference, regarding mean vessel radius, compared to PBS, whilst in 2D there are significant differences between PBS and Captopril, PBS and FLIO, and PBS and Yes (Fig. 5b). We attributed this difference to the elliptical cross-section of retinal vessels. The long axis of the vessels is parallel to the inner limiting membrane, i.e., parallel with respect to the XY plane (Sup. Fig. 6 top). This leads to an overestimation of vessel radius when using 2D MIPs (Sup. Fig. 6 bottom), and better estimation of vessel radius in 3D, since in 3D we quantify the mean radius across the entire contour of the vessel cross-section.

Mean branch length and branching point density followed similar trends to 2D quantifications, yet some of the relative changes, when compared to the PBS group, did not correlate well between 2D and 3D quantifications. For instance, differences in the mean branch length between PBS and AngII and PBS and FLIO are not evident in 3D. Furthermore, our 3D analysis indicated statistically significant differences in branching point density between PBS and Captopril and PBS and AngII, and in the mean branch length between PBS and VEGF, which were not evident in the 2D study (Fig. 5b).

Thus, we demonstrate that 3D quantifications are important to accurately describe vascular morphometrics and that caution must be taken when drawing conclusions from 2D MIPs.

## 4 Discussion

Analysis of 3D vascular networks is an essential step to understand several physiological and pathological processes. In this work, we disclose 3DVascNet, a novel DL-based software for automated segmentation and quantification of 3D retinal vascular networks. Our approach is based on the 3D CycleGAN model which translates 3D images into 3D segmentation masks. Importantly, the proposed method for segmentation only required 2D ground-truth masks to train the 3DVascNet model. The main advantage of the proposed pipeline compared to previous approaches ([4–9]) is that our method can be used to study retinas at any developmental stage since it allows to perform segmentation in 3D. During the later phases (P8 and beyond) the blood vessels in the retina develop a three-layered 3D structure. Analysing in 2D retinas from these stages can lead to incorrect estimation of branching points and branch lengths due to the overlap of vessels from different layers. We have shown that our approach is able to segment 3D blood vessels corresponding to a P20 retina (Sup. Videos 2-4). Thus, the proposed method will clearly be a valuable tool in vascular research.

Even if 3DVascNet produces fully automated high-quality 3D segmentation masks, we strongly recommend the acquisition of microscopy images without the presence of small noisy regions of high intensity, typically caused by the presence of dirt or air-bubbles, as these were incorrectly segmented by our model, compromising the quality of the segmentation masks (Sup. Fig. 7).

We have shown that our method outperforms traditional approaches for segmentation (Otsu thresholding [25] and Chan-Vese [27]). On the one hand, these classical methods are based on image intensities, resulting in over-segmentation in regions with background noise, and under-segmentation in regions with heterogeneous staining. On the other hand, the 3DVascNet performs segmentation based on automatically extracted features from the input images which are learned throughout the training process. Our results show that DL-based segmentation masks represent accurately the vessels in the microscopy images. Furthermore, our results indicate that the proposed approach presents a high generalization capability. Hence, it can be used for accurate segmentation of 3D retinal vessels in microscopy images acquired under various different experimental conditions.

Compared to the method proposed in [18], the proposed approach produces higher-quality 3D segmentation masks. This is likely because, in this work, the 3D segmentation masks are obtained from the 3D images, whereas in [18] the 3D masks are computed based on 2D masks and a set of assumptions, such as the spherical shape of the vessel cross-section and coplanarity of the network. Although 3DVascNet presented a high-generalization capability, it may be necessary to further improve its performance by retraining with new images. Since the source code of 3DVascNet and the datasets are made publicly available, our model can be easily trained with user-specific 3D images and provided 3D masks, which will extend 3DVascNet to other vascular networks. For instance, light-sheet microscopy has been used to image entire vascular networks of the retina ([34, 35]) and the brain ([36]). It would be interesting to test and expand 3DVascNet to segment these types of datasets.

Although the source-codes of most of the described works ([6, 8–10]) are made publicly available, only two works [7, 8] have presented a GUI which can easily be used by researchers without a solid expertise in computer science and programming. To overcome this issue, we present not only 3DVascNet’s source-code which can be easily adapted for the segmentation of other vascular networks, but also a user-friendly graphical interface.

We have validated the performance of 3DVascNet by comparing its output to the quantification of 4 vascular morphometric parameters based on ground-truth segmentation masks. Our results indicated that there are no statistically significant differences between the vascular morphometric parameters obtained from the manual masks and 2D projections of 3DVascNet’s masks.

Also, there was a high linear correlation between manual and 3DVascNet-based computation of vascular morphometric parameters (*R*^2^ *≥* 0.87). Thus, the quantitative analyses yielded similar results compared to manual segmentation. However, 2D analysis of retinal vasculature has several limitations. For instance, 2D studies may be misleading due to false positive branches and branching points resulting from crossings of various vessels along the z-axis. This is due to the fact that the 2D quantifications do not take into consideration the true geometry of the vessels. Hence, we also computed the 3D features from the 3DVascNet’s masks and corresponding skeletons. Although the ranges of branch lengths across different intervention groups are similar in 3D and 2D, there is a notable disparity in the quantification of the vessel radius between 2D and 3D. These differences are likely explained by the assumption that vessels cross-sections are circular, whilst, in reality, they are oval, leading to an overestimation of vessel radii in 2D. Thus, the 3D quantifications obtained with the proposed approach are superior in comparison to the traditional 2D analysis of retinal blood vessels. This is due to their ability to capture the volumetric information providing a more realistic quantification of the 3D blood vessels. To conclude, the differences between the 2D and 3D quantifications enhance the need of undertaking 3D analysis of the vasculature instead of studying its 2D projections.

To summarize, in this work, we have presented 3DVascNet, a software for segmentation and quantification of blood vessels in 3D. 3DVascNet is fully automated, thus it will be used to perform large-scale analysis in 3D, which will be of great importance to answer key open questions in vascular biology.

## Disclosures

### Ethics approval

As stated in [6], animal experimentation was carried out in compliance with EU Directive 86/609/EEC and Recommendation 2007/526/EC regarding the protection of animals used for experimental and other scientific purposes. Animal procedures were performed under supervision by the IMM-JLA Animal Ethics Committee (ORBEA), under the project licenses AWB 2015 10 CF Polaridade and AWB 2021 02 CF Vascular, and approved by the Portuguese Animal Ethics Committee regulatory body (DGAV), project licenses 0421/000/000/2016 and 0421/000/000/2021.

### Consent to participate

Not applicable

### Consent for publication

Not applicable

### Availability of data and materials

The datasets used in this work will be made available at https://huggingface.co/datasets/Hemaxi/ 3DVesselSegmentation.

### Code Availability

The code will be made available at https://github.com/ HemaxiN/3DVascNet.

### Authors’ contributions

Conceptualization of this study: Hemaxi Narotamo, Margarida Silveira and Cláudio Areias Franco.

Methodology: Hemaxi Narotamo, Margarida Silveira and Cláudio Areias Franco.

Investigation: Hemaxi Narotamo, Margarida Silveira and Cláudio Areias Franco.

Software: Hemaxi Narotamo.

Validation: Hemaxi Narotamo, Margarida Silveira and Cláudio Areias Franco.

Visualization: Hemaxi Narotamo, Margarida Silveira and Cláudio Areias Franco.

Supervision: Margarida Silveira and Cláudio Areias Franco. Writing - Original draft preparation: Hemaxi Narotamo.

Writing - review and editing: Hemaxi Narotamo, Margarida Silveira and Cláudio Areias Franco.

## Acknowledgments

We thank Rui Benedito (Centro Nacional de Investigaciones Cardiovasculares Carlos III), and Lena Claesson-Welsh and Jin Yi (Uppsala University) for providing images or retinas used in this study. We thank Miguel Bernabeu (Edinburgh University) for insights and comments on the usage of PolNet. We are grateful to Ana Figueiredo, Andreia Pena, Daniela Ramalho, Lenka Misiková and Marta Saraiva for their valuable suggestions on the GUI. We would like to thank Pedro Barbacena, Marie Ouarńe, Ana Figueiredo, Andreia Pena and Daniela Ramalho for providing the microscopy images used in this study. We thank Pedro Barbacena, Andreia Pena, Daniela Ramalho for manually annotating the 2D ground-truth masks used in this work.

## Footnotes

1 Jason Brownlee, “How to develop a CycleGAN for image-to-image translation with Keras”, https://machinelearningmastery.com/cyclegan-tutorial-with-keras/, 2019.

2 3DVascNet is currently available for Windows.

3 Although the ROIs are defined in 2D, the final ROIs are 3D, which are computed as explained in Subsection 2.2.3. That is, the final 3D ROIs are the intersection of the 3D convex hull with the 2D ROIs.

---

**Supplementary Fig. 1:**
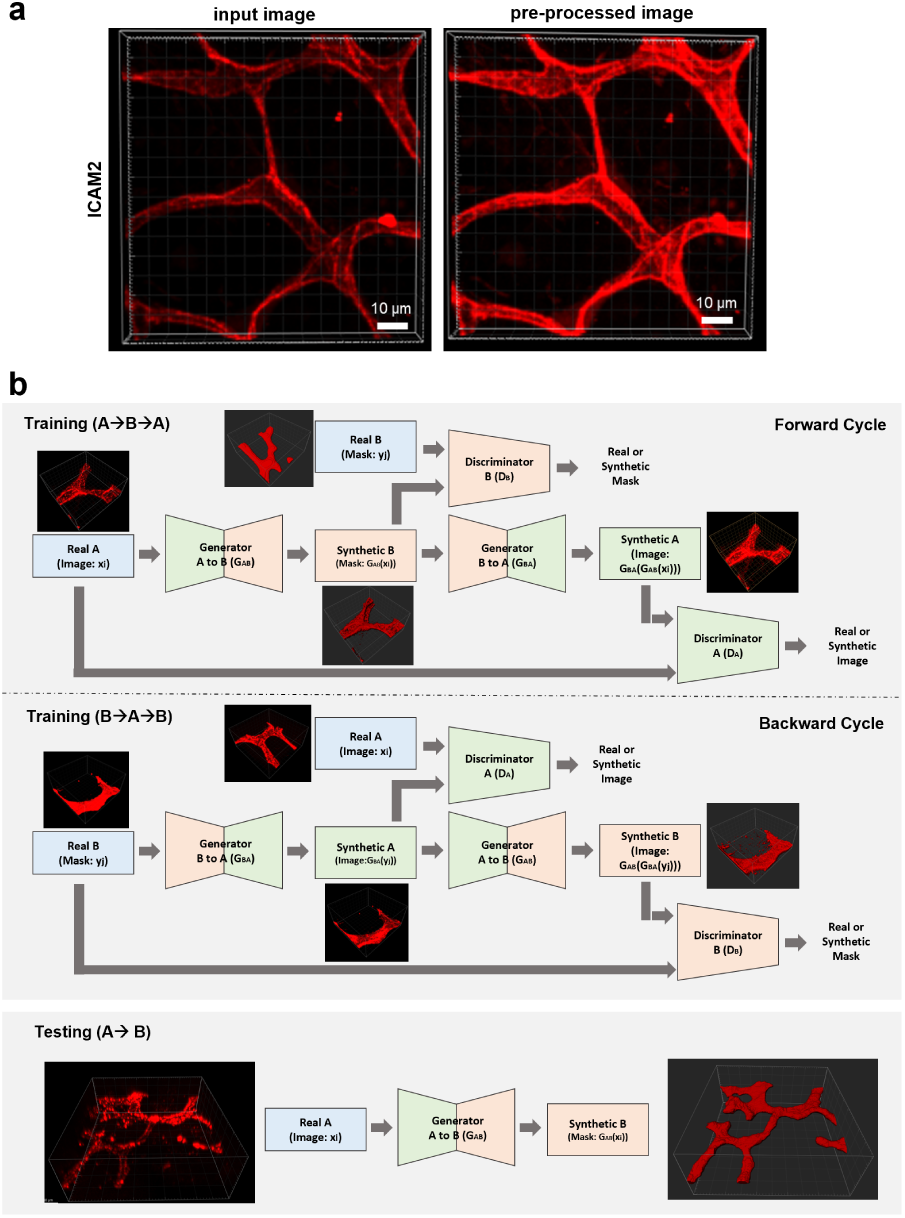
3D CycleGAN for 3D retinal vessel segmentation in microscopy images. **a** Representative image showcasing the effects of percentile normalization. Left image, patch extracted from the original image containing retinal blood vessels; right image, the corresponding patch extracted from the pre-processed image. **b** 3D CycleGAN for 3D retinal vessel segmentation in microscopy images. Top: Illustration of the adversarial training (*G_AB_ ↔ D_B_* and *G_BA_ ↔ D_A_*) and of the two training paths of 3D CycleGAN: forward and backward cycles to compute the forward and backward cycle-consistency losses *x → G_AB_*(*x*) *→ G_BA_*(*G_AB_*(*x*)) *≈ x* (on the top) and *y → G_BA_*(*y*) *→ G_AB_*(*G_BA_*(*y*)) *≈ y* (on the bottom), respectively; Bottom: Illustration of the test step: the generator A to B receives as input a patch extracted from the microscopy image and outputs the corresponding segmentation mask.

**Supplementary Fig. 2:**
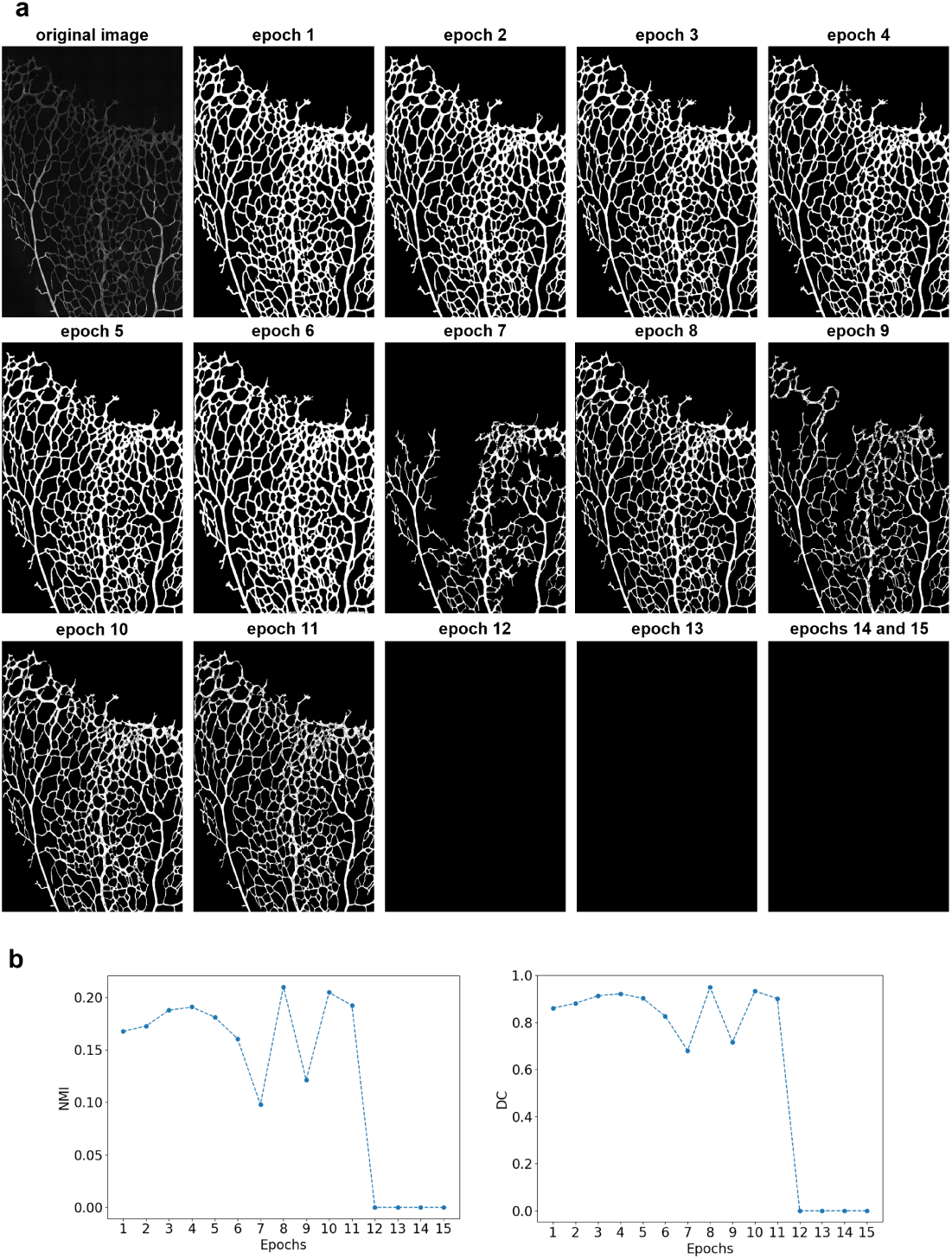
Effects of different epochs in the quality of 3D CycleGAN image segmentation. **a** 2D projections of a training image and respective CycleGAN’s outputs over the 15 training epochs. To select the best epoch of the 3D CycleGAN model we not only visually inspected the output of the model for the training images, but also considered the NMI metric. We visually inspected the segmentation masks obtained for the training images corresponding to the epochs that present the highest values of NMI: epochs 4, 8 and 10 that present NMI values of 0.19, 0.21 and 0.20, respectively. Visual inspection of the segmentation masks for epochs 4, 8 and 10 revealed that the masks obtained for epochs 8 and 10 present a lot of discontinuities and holes, thus we selected the model saved at epoch 4 to evaluate its performance on the test images. **b** Plots of the NMI and DC values over the 15 training epochs. We have shown that the NMI metric is consistent with the supervised DC evaluation metric.

**Supplementary Fig. 3:**
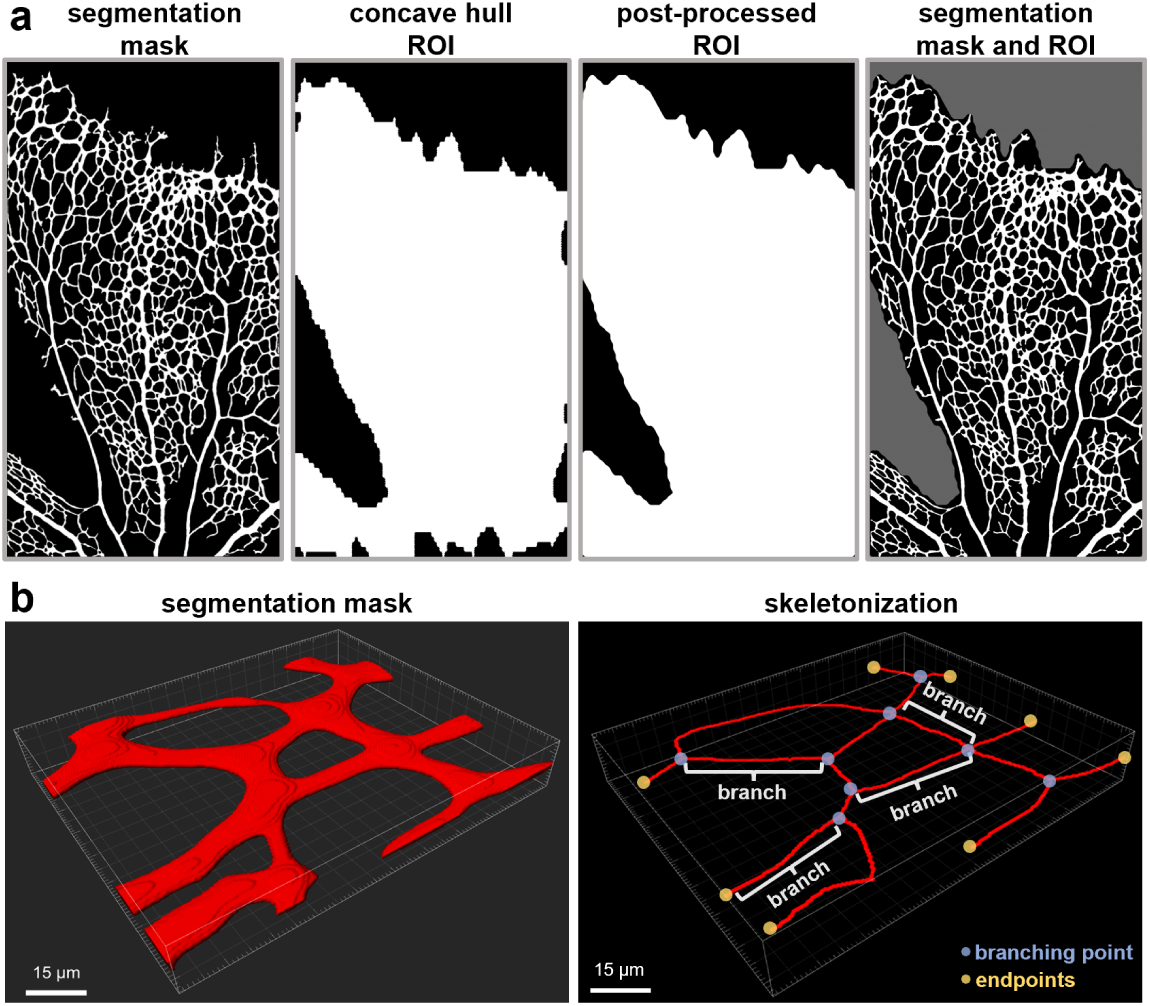
Quantification of vascular morphometrics using 3DVascNet. a Representative images demonstrating the automatic selection of the region of interest (ROI) based on the concave hull method: from left to right: binary segmentation mask of vessels; corresponding ROI computed based on the method proposed in [22]; corresponding post-processed ROI (inclusion in the ROI of small background regions, and median filtering); and representation of the vessels’ segmentation mask (in white), and region excluded from the analysis (in gray). **b** Example of automated skeletonization and parameter extraction. Left: 3D patch extracted from a segmentation mask; Right: corresponding skeleton represented in red, some branches are identified in the skeleton in white, the branching points are represented in blue, and the endpoints are shown in yellow.

**Supplementary Fig. 4:**
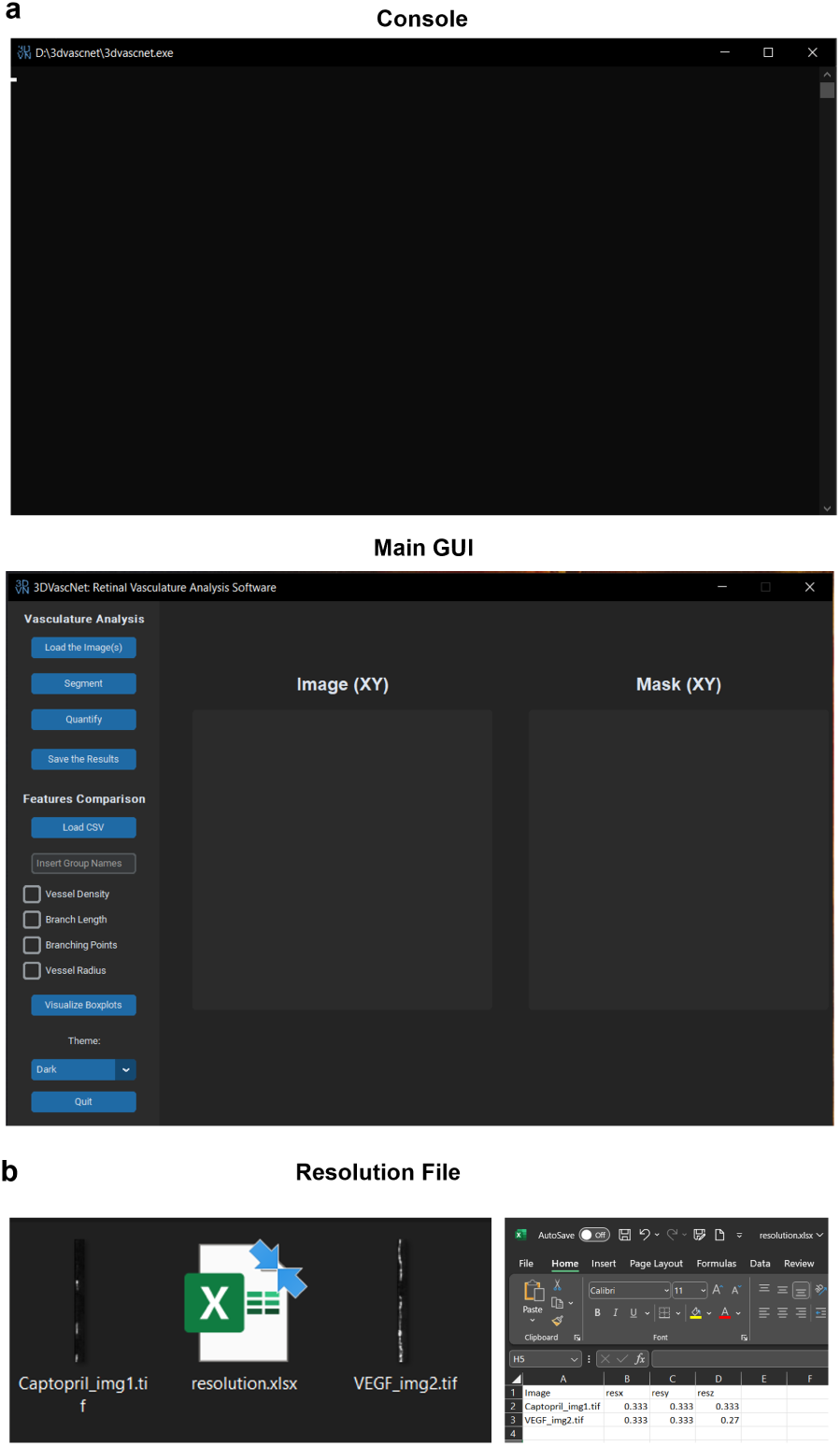
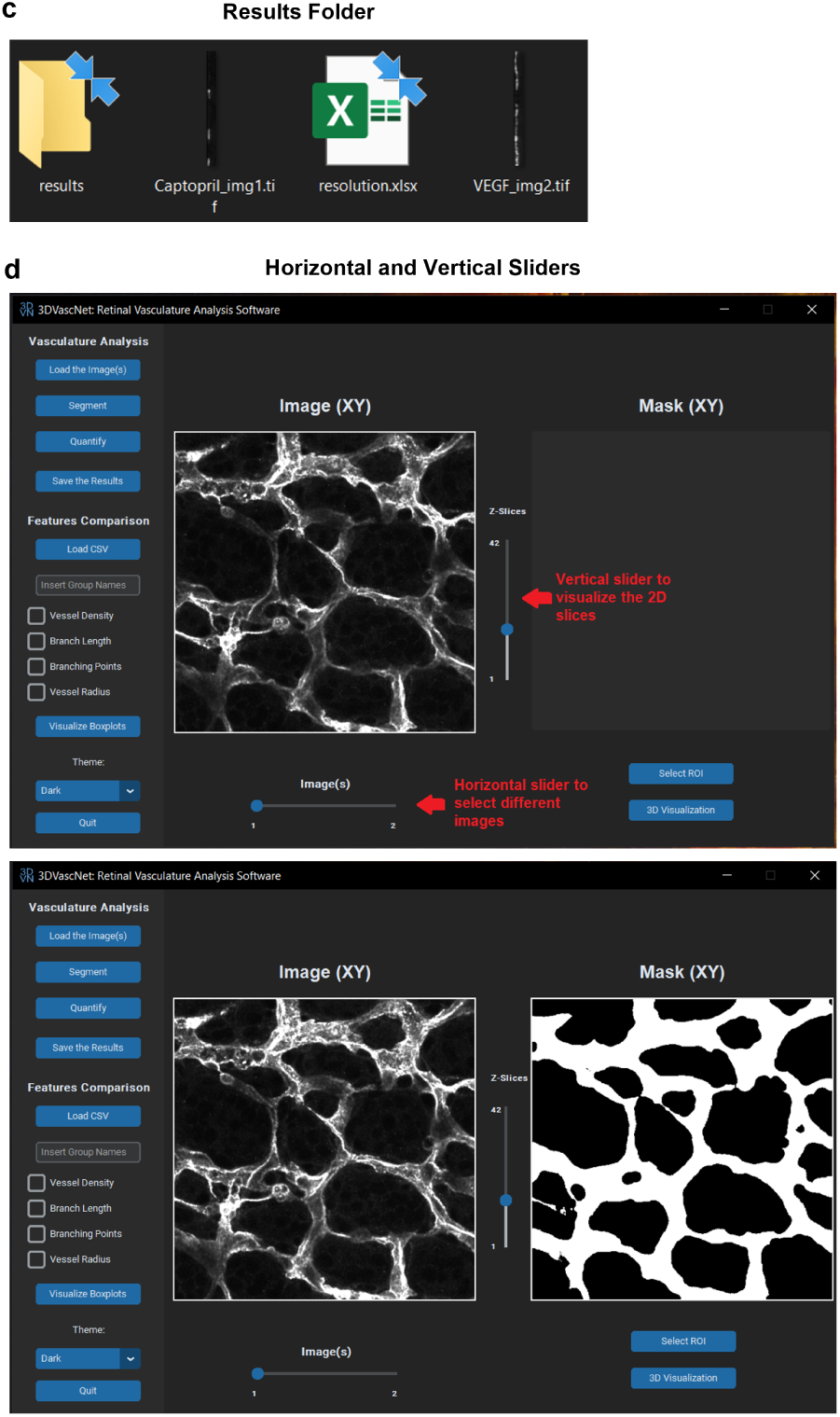
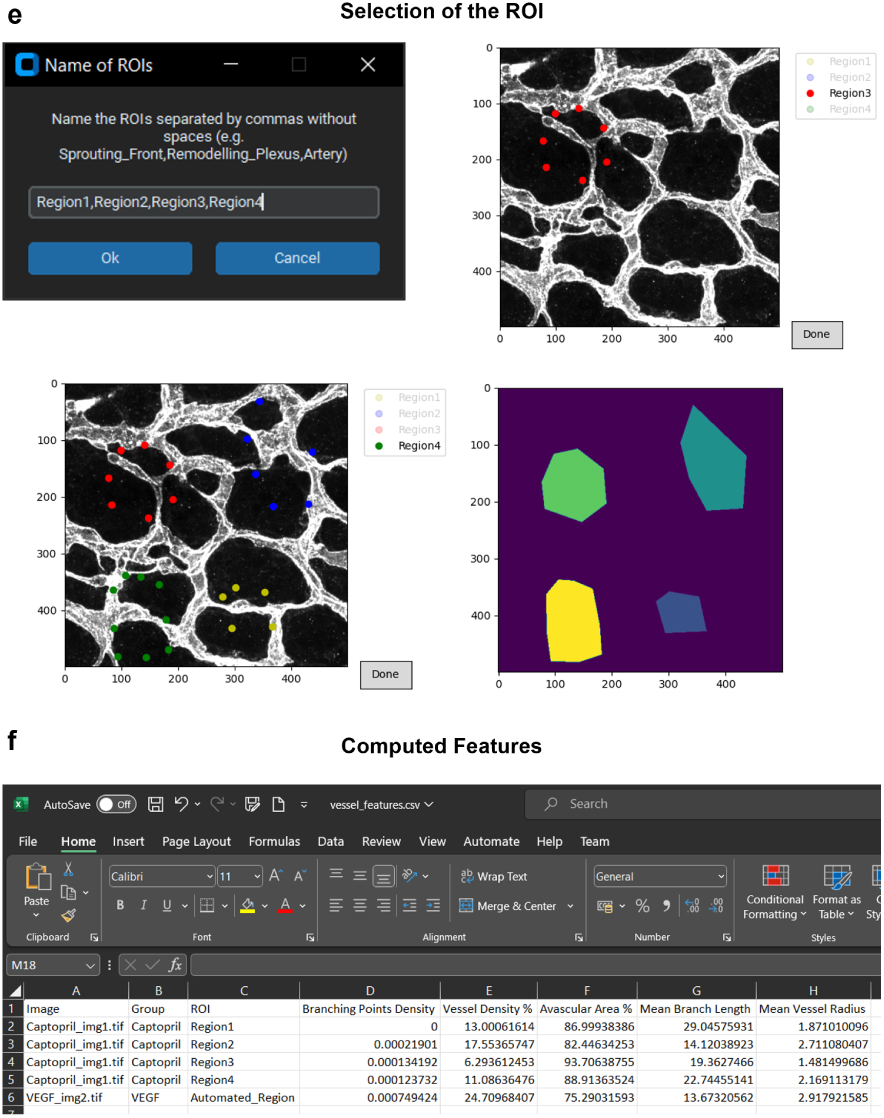
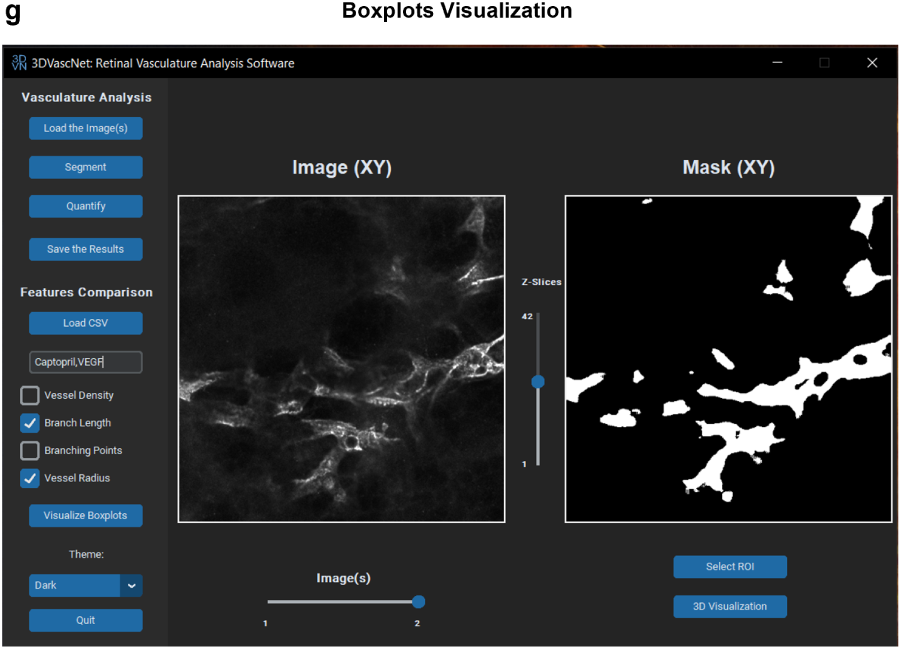
3DVascNet’s GUI. **a** 3DVascNet application running on a machine, it has two components: a console that displays outputs and errors, and the main GUI. **b** Files in the directory selected upon clicking on the “Load Image(s)” button: two .tif images and a resolution.xlsx file, and content of the resolution.xlsx file, it should contain, for each image in the directory, the values of the voxel’s physical dimensions along the three dimensions. **c** The automatically created results folder in the selected directory. **d** Illustration of the horizontal and vertical sliders that can be used to select different images and slices of each image, respectively. **e** Manual ROI selection process: specification of the ROIs names, selection of region 3, completed selection of the 4 ROIs, manually defined ROIs with different colors. **f** Vessel features.csv file containing the computed features. For “Captopril img1.tif” image the features were computed for the four regions that were manually defined (**e**), and for “VEGF img2.tif” image the features were computed based on the automatically computed 3D concave hull. **g** Group names inserted in the “Insert Group Names” box and selection of the Branch Length and Vessel Radius features. 3DVascNet application running on a machine, it has two components: a console that displays outputs and errors, and the main GUI. **b** Files in the directory selected upon clicking on the “Load Image(s)” button: two .tif images and a resolution.xlsx file, and content of the resolution.xlsx file, it should contain, for each image in the directory, the values of the voxel’s physical dimensions along the three dimensions. **c** The automatically created results folder in the selected directory. **d** Illustration of the horizontal and vertical sliders that can be used to select different images and slices of each image, respectively. **e** Manual ROI selection process: specification of the ROIs names, selection of region 3, completed selection of the 4 ROIs, manually defined ROIs with different colors. **f** Vessel features.csv file containing the computed features. For “Captopril img1.tif” image the features were computed for the four regions that were manually defined (**e**), and for “VEGF img2.tif” image the features were computed based on the automatically computed 3D concave hull. **g** Group names inserted in the “Insert Group Names” box and selection of the Branch Length and Vessel Radius features. 3DVascNet application running on a machine, it has two components: a console that displays outputs and errors, and the main GUI. **b** Files in the directory selected upon clicking on the “Load Image(s)” button: two .tif images and a resolution.xlsx file, and content of the resolution.xlsx file, it should contain, for each image in the directory, the values of the voxel’s physical dimensions along the three dimensions. **c** The automatically created results folder in the selected directory. **d** Illustration of the horizontal and vertical sliders that can be used to select different images and slices of each image, respectively. **e** Manual ROI selection process: specification of the ROIs names, selection of region 3, completed selection of the 4 ROIs, manually defined ROIs with different colors. **f** Vessel features.csv file containing the computed features. For “Captopril img1.tif” image the features were computed for the four regions that were manually defined (**e**), and for “VEGF img2.tif” image the features were computed based on the automatically computed 3D concave hull. **g** Group names inserted in the “Insert Group Names” box and selection of the Branch Length and Vessel Radius features. 3DVascNet appli-cation running on a machine, it has two components: a console that displays outputs and errors, and the main GUI. **b** Files in the directory selected upon clicking on the “Load Image(s)” button: two .tif images and a resolution.xlsx file, and content of the resolution.xlsx file, it should contain, for each image in the directory, the values of the voxel’s physical dimensions along the three dimensions. **c** The automatically created results folder in the selected directory. **d** Illustration of the horizontal and vertical sliders that can be used to select different images and slices of each image, respectively. **e** Manual ROI selection process: specification of the ROIs names, selection of region 3, completed selection of the 4 ROIs, manually defined ROIs with different colors. **f** Vessel features.csv file containing the computed features. For “Captopril img1.tif” image the features were computed for the four regions that were manually defined (**e**), and for “VEGF img2.tif” image the features were computed based on the automatically computed 3D concave hull. **g** Group names inserted in the “Insert Group Names” box and selection of the Branch Length and Vessel Radius features.

**Supplementary Fig. 5:**
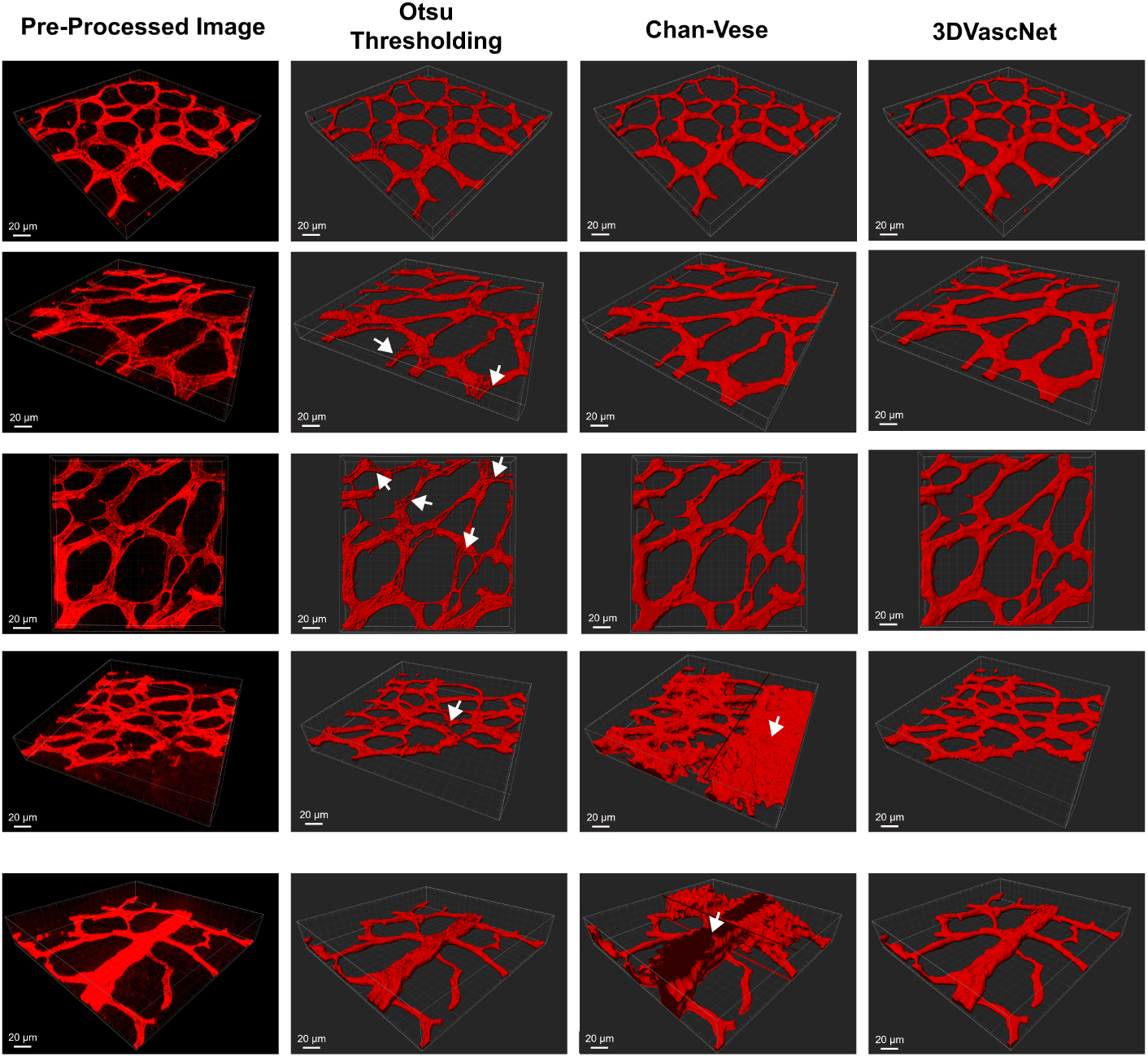
3DVascNet’s segmentation performance comparison with two classical approaches. Examples of 3D patches extracted from the pre-processed images (first column), and the corresponding segmentation masks obtained with Otsu thresholding (second column), Chan-Vese (third column) and 3DVascNet (fourth column). The white arrows indicate regions poorly segmented by Otsu due to heterogeneous staining, and background noise segmented by Chan-Vese.

**Supplementary Fig. 6:**
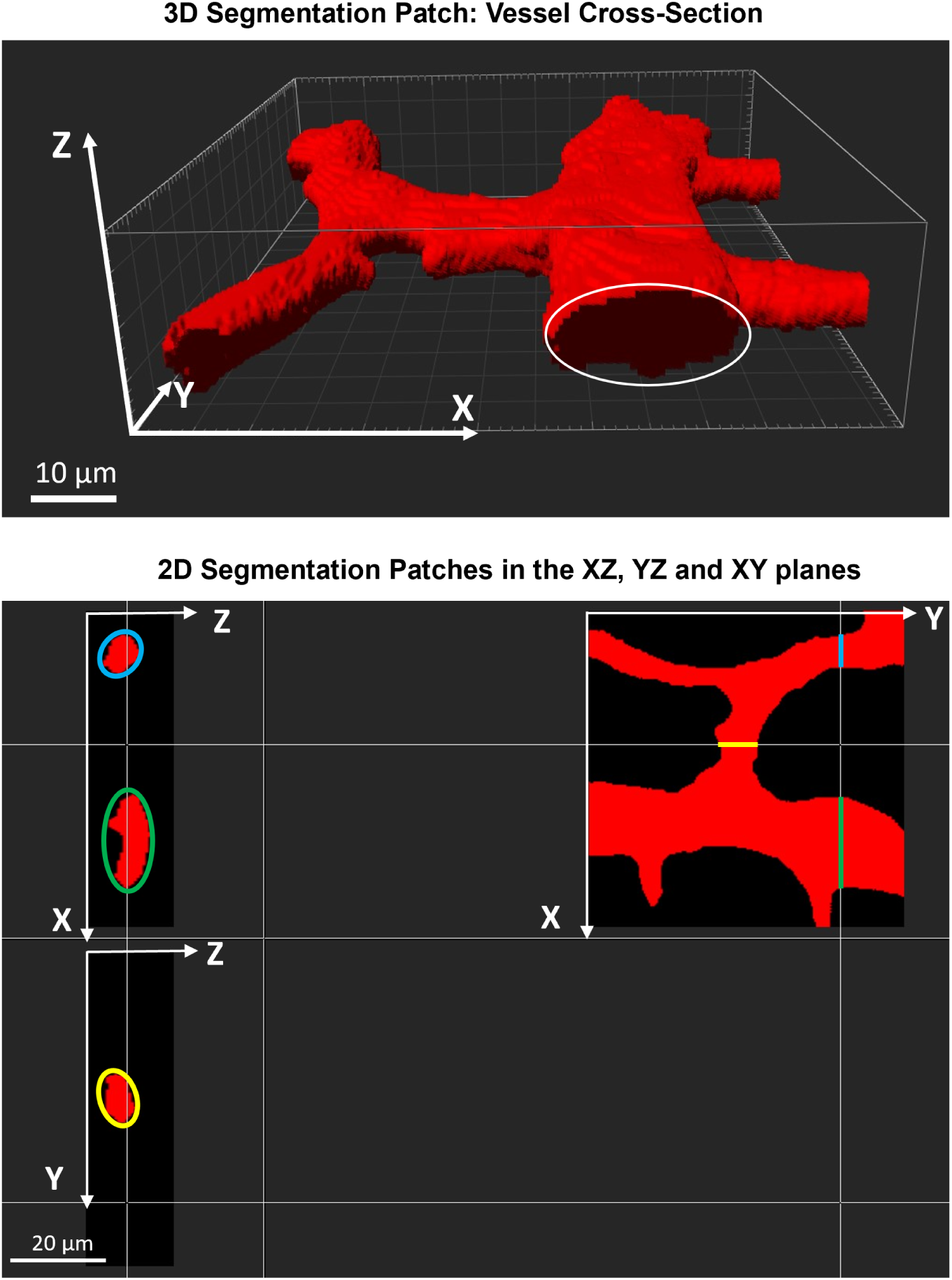
3D Vessels’ Elliptical Cross-Section. Top: 3D patch obtained from a 3DVascNet’s segmentation mask illustrating the vessels aligned with the XY plane, and the elliptical cross-section of the vessel aligned with the Y direction (indicated with a white ellipse). Bottom: Corresponding 2D sections in the XZ, YZ and XY planes. The radius in 2D is computed based on the projection of the mask on the XY plane leading to an overestimation of the radius.

**Supplementary Fig. 7:**
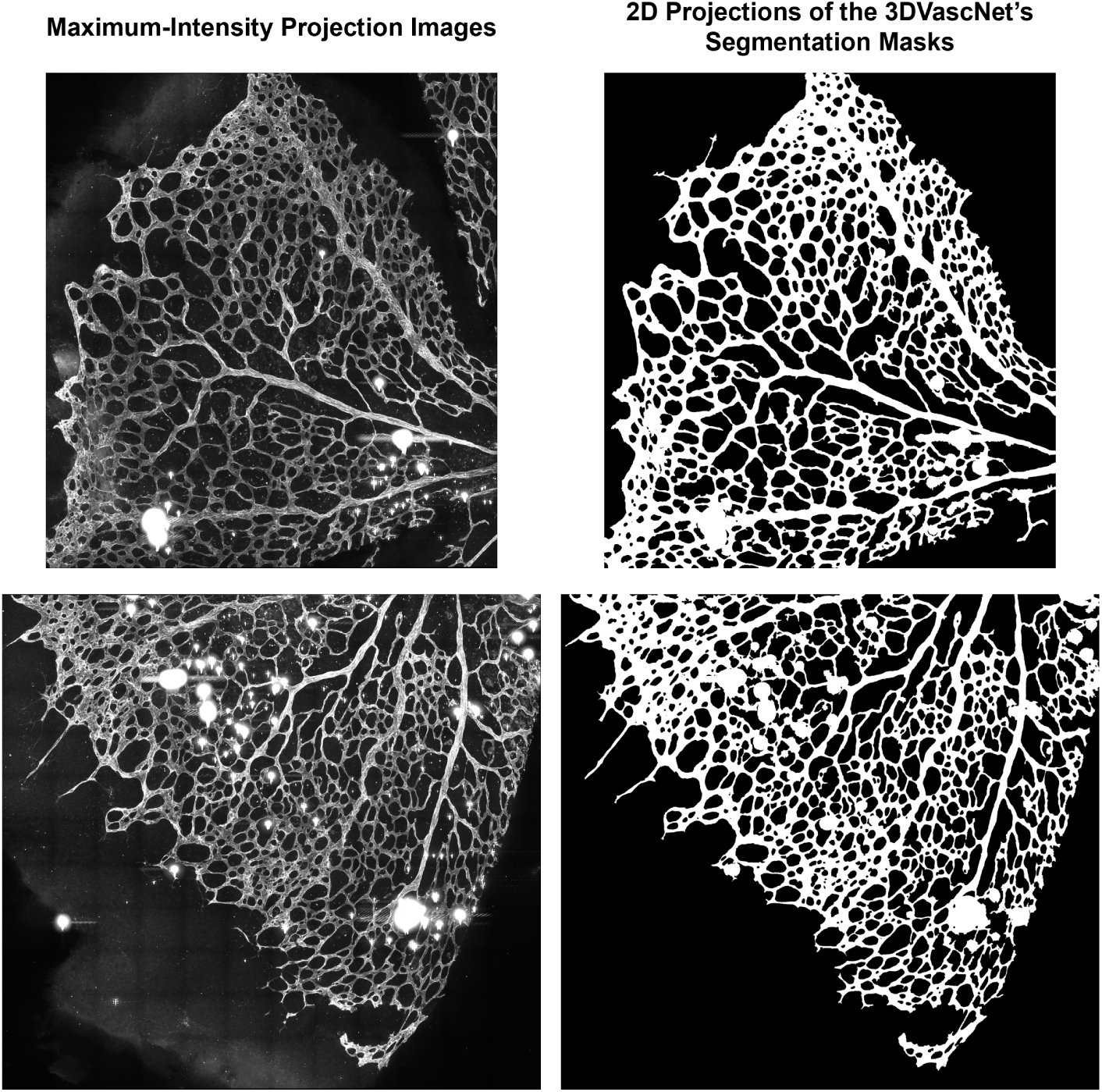
Effect of noise in the 3DVascNet’s segmentation output. Analysis of noisy images from the VEGF group: On the left, MIPs of two images containing noise (small blobs with high intensity); on the right, corresponding 2D projections of the 3D segmentation masks obtained with 3DVascNet.

**Supplementary Table 1:** Sizes of the test images from datasets 2, 3 and 4. The retinas with the prefix YES were reported in [37].

| | Image | Size ($X \times Y \times Z$) |
| --- | --- | --- |
| **Dataset 2** | Captopril_ret1 | $4212 \times 5427 \times 35$ |
| | Captopril_ret2 | $4194 \times 5013 \times 34$ |
| | Captopril_ret3 | $4599 \times 3789 \times 36$ |
| | Captopril_ret4 | $3206 \times 3972 \times 37$ |
| | Captopril_ret5 | $3204 \times 4365 \times 54$ |
| | Captopril_ret6 | $3600 \times 4365 \times 60$ |
| | RGDS_ret1 | $3553 \times 4000 \times 67$ |
| | RGDS_ret2 | $2258 \times 3997 \times 42$ |
| | RGDS_ret3 | $3555 \times 4428 \times 48$ |
| **Dataset 3** | FLIJ_ret1 | $4590 \times 2160 \times 27$ |
| | FLIJ_ret2 | $3384 \times 5427 \times 35$ |
| | FLIJ_ret3 | $2709 \times 5733 \times 41$ |
| | FLIJ_ret4 | $3402 \times 5445 \times 35$ |
| | FLIO_ret1 | $3388 \times 2969 \times 32$ |
| | FLIO_ret2 | $3783 \times 3375 \times 43$ |
| | FLIO_ret3 | $3378 \times 3787 \times 28$ |
| | FLIO_ret4 | $5004 \times 3798 \times 27$ |
| | FLIO_ret5 | $4428 \times 2700 \times 40$ |
| | FLIO_ret6 | $4203 \times 4212 \times 31$ |
| | FLIO_ret7 | $4608 \times 5022 \times 44$ |
| | FLIO_ret8 | $5004 \times 3798 \times 38$ |
| **Dataset 4** | YES_ret1 | $5632 \times 6656 \times 36$ |
| | YES_ret2 | $4608 \times 5632 \times 36$ |
| | YES_ret3 | $4608 \times 4608 \times 28$ |
| | YES_ret4 | $4608 \times 4608 \times 32$ |
| | YES_ret5 | $4608 \times 4608 \times 24$ |
| | YES_ret6 | $4608 \times 5632 \times 20$ |
| | YES_ret7 | $4608 \times 5632 \times 24$ |

## Supplementary Information I Downloading and Running 3DVascNet

### 1 Downloading and Launching the Application

- Download the 3dvascnet.zip available at https://github.com/HemaxiN/ 3DVascNet^2^;
- Extract the files in your working directory, this will create a folder named 3dvascnet with the application files.
- To start the application, double-click on the 3dvascnet.exe file, two windows will open: a console that will display output and error messages, and the main GUI (Sup. Fig. 4a).

### 2 Loading the Images

- Click on the “Load Image(s)” button and select the directory containing the images that will be analyzed. Currently, our software supports .tif and .czi files, each image must have three-dimensions (X,Y,Z), and one channel representing the blood vessels. The images’ names should have a prefix denoting its group. This is important to later visualize the distributions of the vascular features by grouping images belonging to the same group.
- Apart from the images, this directory should also contain a resolution.xlsx file (Sup. Fig. 4b left) with the information, for each image, about the voxel’s physical size (in *µm*) along the x, y and z axes (Sup. Fig. 4b right). 3DVascNet will automatically read this file, or throw an error message if the file is not present in the selected directory.
- 3DVascNet will automatically create a results folder in the selected directory where all results will be saved (Sup. Fig. 4c).
- (OPTIONAL) To confirm that the images were properly loaded, use the horizontal slider to visualize different images, and the vertical slider to see each 2D slice across the depth dimension (Sup. Fig. 4d). Herein, we used two small images to explain the method, users can analyze entire retinas with our software.

### 3 Performing Segmentation

- Click on the “Segment” button, this will perform segmentation of all the images based on the proposed 3D CycleGAN model. This step is computationally demanding, the segmentation time will highly depend on the number and size of the images and the specifications of the machine. It is recommended to use a machine with an Nvidia GPU. Output messages in the console will allow to monitor the segmentation process.
- The segmentation mask of each image will be saved in the results folder as a .tif file.
- The sliders can be used again to visualize the images and corresponding segmentation masks (Sup. Fig. 4d).

### 4 Performing Quantification

- (OPTIONAL) Select an image using the horizonal slider for which several ROIs need to be defined, then click on the “Select ROI” button, insert the names of the ROIs to be annotated. The names should be separated by commas and should not have spaces between them (Sup. Fig. 4e). Thereafter, manually select each ROI after clicking on the ROI name on the right. Using the left mouse button, add points to define the ROI for the selected region name (Sup. Fig. 4e). The region can be corrected clicking the right mouse button while placed on top of a previously defined point. After selecting the first ROI, click on another region name from the list on the right and add the points to define the ROI. When all ROIs are defined click on the “Done” button (Sup. Fig. 4e) ^3^.
- Click on the “Quantify” button to perform 3D vasculature quantification. Output messages in the console will allow to monitor the quantification process. 3D skeletons will be automatically saved in the results folder, as .tif files with the “Skeleton “ prefix in their names.
- Once finished, click on the “Save the Results” button, which will save a vessel features.csv file in the results folder containing the computed features (Sup. Fig. 4f). A previously generated .csv file can be loaded at any time using the “Load CSV” button.

### 5 Comparison of Retinal Vasculature Features

- Type the group names separated by commas and without spaces between them in the “Insert Group Names” box, for this simple example the following should be typed: **Captopril,VEGF**. Then select the features (Vessel Density, Branch Length, Branching Points and/or Vessel Radius) (Sup. Fig. 4g), and click on the “Visualize Boxplots” button to compare retinal vasculature distributions for different groups.

### 6 Close the Application

- To stop the application click on the “Quit” button or close the GUI and console windows.

## Supplementary Videos

**Supplementary Video 1**: https://drive.google.com/file/d/1kGwHIY0 2gW1dPn Hg60 DG8VB4YJDF9/view?usp=drive link **Automated definition of the 3D ROI.** Left: Slices along the z-axis extracted from the 3D segmentation mask obtained with the 3D CycleGAN model. Right: Slices along the z-axis extracted from the 3D ROI that corresponds to the intersection of each slice of the 3D convex hull with the 2D concave hull computed from the 3D mask’s projection.

**Supplementary Video 2**: https://drive.google.com/file/d/ 1TJGmjIBAdQewF5D8Sew6FJK8SSsftxhz/view **P20 Retina: Image.** Representative patch extracted from an image corresponding to a P20 retina to illustrate the three-dimensionality of the network.

**Supplementary Video 3**: https://drive.google.com/file/d/ 149OXYiSPMA42rWJWpONi5EaqENnxro28/view?usp=sharing **P20 Retina: Mask.** Segmentation mask of the patch shown in Sup. Video 2, obtained with 3DVascNet.

**Supplementary Video 4**: https://drive.google.com/file/d/ 1IOPR4jLhNA50ApLp43PVl-nHluAQGCYq/view?usp=sharing **P20 Retina: Image and Mask Overlap.** Visualization of the overlap between the image (red) and the segmentation mask (green) shown in Sup. Videos 2 and 3, respectively

